# Distinct Myeloid Derived Suppressor Cell Populations Promote Tumor Aggression in Glioblastoma

**DOI:** 10.1101/2023.03.26.534192

**Authors:** Christina Jackson, Christopher Cherry, Sadhana Bom, Arbor G. Dykema, Elizabeth Thompson, Ming Zheng, Zhicheng Ji, Wenpin Hou, Runzhe Li, Hao Zhang, John Choi, Fausto Rodriguez, Jon Weingart, Srinivasan Yegnasubramanian, Michael Lim, Chetan Bettegowda, Jonathan Powell, Jennifer Eliesseff, Hongkai Ji, Drew Pardoll

## Abstract

The diversity of genetic programs and cellular plasticity of glioma-associated myeloid cells, and thus their contribution to tumor growth and immune evasion, is poorly understood. We performed single cell RNA-sequencing of immune and tumor cells from 33 glioma patients of varying tumor grades. We identified two populations characteristic of myeloid derived suppressor cells (MDSC), unique to glioblastoma (GBM) and absent in grades II and III tumors: i) an early progenitor population (E-MDSC) characterized by strong upregulation of multiple catabolic, anabolic, oxidative stress, and hypoxia pathways typically observed within tumor cells themselves, and ii) a monocytic MDSC (M-MDSC) population. The E-MDSCs geographically co-localize with a subset of highly metabolic glioma stem-like tumor cells with a mesenchymal program in the pseudopalisading region, a pathognomonic feature of GBMs associated with poor prognosis. Ligand-receptor interaction analysis revealed symbiotic cross-talk between the stemlike tumor cells and E-MDSCs in GBM, whereby glioma stem cells produce chemokines attracting E-MDSCs, which in turn produce growth and survival factors for the tumor cells. Our large-scale single-cell analysis elucidated unique MDSC populations as key facilitators of GBM progression and mediators of tumor immunosuppression, suggesting that targeting these specific myeloid compartments, including their metabolic programs, may be a promising therapeutic intervention in this deadly cancer.

**One-Sentence Summary:** Aggressive glioblastoma harbors two unique myeloid populations capable of promoting stem-like properties of tumor cells and suppressing T cell function in the tumor microenvironment.

## Main Text

Checkpoint blockade immunotherapy targeting the T cell checkpoints CTLA-4, PD-1/L1 and recently LAG3 (*1–3*) has shifted the treatment standard of over 18 solid tumors. However, the benefit of these immune based therapies in glioblastoma multiforme (GBM) has thus far been disappointing. GBM, the most lethal high grade form of glioma, is a quintessential example of an immunologically “cold tumor” with relatively few tumor infiltrating lymphocytes, thus greatly limiting the effectiveness of T cell targeted therapies. Nonetheless, true neoantigen-specific T cells have been demonstrated in GBM (*4*). Taken together, these findings suggest that other cells and signals in the GBM microenvironment could suppress the function of tumor-specific T cells. Tumor infiltrating myeloid cells are an important component of the GBM microenvironment (TME) constituting >70% of tumor infiltrating immune cells in GBM (*5*). However, the precise composition and phenotypes of glioma associated myeloid cells (GAMs), their role in shaping and interacting with other components of the GBM TME, and their tractability as targets for therapeutic intervention are poorly understood.

In order to better characterize the cellular components of primary brain cancer, we performed single-cell RNA sequencing (scRNA-seq) on a cohort of gliomas (n=33) spanning low to high grade tumors, focusing on populations uniquely acquired in grade IV GBMs. Surveying the transcriptomes of >750,000 immune cells and >350,000 tumor and associated stromal cells in these samples, we found a diverse landscape of myeloid-lineage cells in gliomas. Two immature bone marrow derived myeloid (BMDM) populations, characteristic of a recently described early progenitor and a monocytic myeloid derived suppressor cell (E- and M-MDSC) population, were almost exclusively present in aggressive grade IV GBM (*6*). Both populations of MDSCs demonstrated strong upregulation of multiple catabolic and anabolic pathways with associated induction of hypoxia and stress response genes. These MDSC populations co-localized with a stem-like tumor cell population in the pseudopalisading region, a pathognomonic feature of GBMs and associated with poor prognosis. Receptor-ligand interaction mapping demonstrates crosstalk between these MDSCs and tumor cells, whereby the tumor cells produce multiple chemokines that would attract the MDSCs and the MDSCs produce factors promoting stem-like features of the co-localized cancer cells. Our findings identify new potential therapeutic targets for improving the efficacy of immune-based therapies in GBM. This large-scale single cell atlas of glioma tumor and immune cells can serve as the foundation for continued exploration of glioma tumor-immune landscapes to further guide T cell independent immunotherapy strategies.

## Results

### ScRNA-seq reveals diverse and distinct landscape of myeloid cells in glioma compared to normal brain

Initially, to characterize the diverse landscape of GAMs in glioma and healthy control patients, we performed scRNA-seq on freshly isolated fluorescence-activated cell sorting (FACS)-purified CD45+CD3-immune cells isolated from 21 resected GBM (grade IV), 7 grade II gliomas (6 oligodendroglioma, one astrocytoma), 5 anaplastic astrocytoma (grade III), and 5 non-malignant brain tissue (Figure 1A, Table S1). In total 240,183 CD3-immune cells passed quality control and were carried forward for analysis (Figure S1A, B). Unbiased clustering and batch effect correction algorithms applied to the total combined transcriptomes from all the analyzed tumors and normal brains subseted CD45+CD3-immune cells into 18 clusters based on global gene expression patterns (Figure S2A). Canonical marker genes were used to identify major cell populations, myeloid cells with low expression of HLA-DR are characteristic of myeloid derived suppressor cells (MDSCs) (Figure 1B). Focusing specifically on myeloid-lineage cells, we identified 14 clusters of myeloid cells including 5 clusters of microglia and 9 clusters of bone marrow-derived myeloid cells (BMDMs) (Figure 1C). Using differentially expressed genes by each cluster compared to other myeloid clusters, we annotated the individual clusters into distinct cell types (Figure 1D-E, Figure S2B, Table S2). Our single-cell profiling demonstrates a diverse and complex composition of microglial and BMDM populations across non-malignant and tumor-bearing CNS compartments that significantly expands upon the traditional view of M1 and M2 tumor associated macrophages in gliomas.

**Figure 1:**
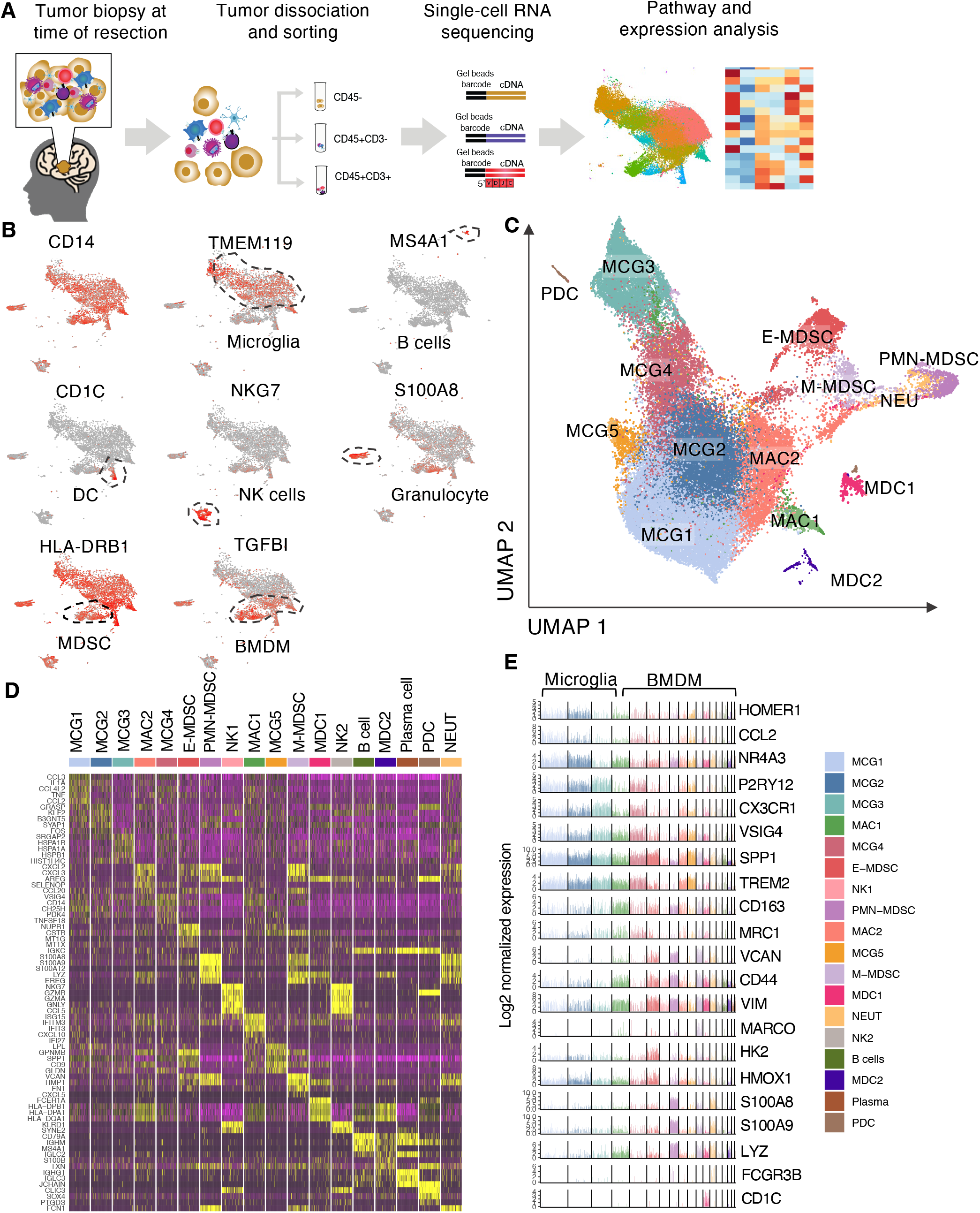
Profiling glioma associated myeloid cells by s cRNA-seq reveals diverse landscape. (A) Schematic graph showing the experimental design of scRNA-seq analysis of tumor and immune cells in gliomas. scRNA-seq was performed on CD45-, CD45+CD3-, and scRNA-seq-TC R-seq was performed on T cells isolated from resected glioblastoma (*n*=21), anaplastic astrocytoma (*n*=5), oligodendroglioma (*n*=6), astrocytoma (*n*=1), and non-neoplastic cortex (*n*=6). (B) UMAP projection of expression of canonical genes for microglia, bone marrow-derived monocytes (BMDM), granulocytes, B cells, NK cells, dendritic cells, and lack of expression of MHCII in myeloid derived suppressor cells (MDSC) (C) UMAP projection of the expression profile of myeloid-lineage cells in gliomas demonstrating 14 cluster of myeloid lineage cells each delineated by color code. (D) Z score normalized heatmap of top 10 differentially expressed genes with highest log fold-change from each cluster. (E) Bar plots showing gene expression levels of select representative genes across microglia and BMDM clusters.

### Distinct myeloid derived suppressor cell populations uniquely present in grade IV glioblastomas

Based on evidence from murine models that specific myeloid populations in the TME can affect the growth and phenotype of tumor cells as well as anti-tumor immunity, we first investigated whether there were differences in distribution of glioma infiltrating myeloid (GAM) populations as a function of tumor grade. While the distribution of GAMs varied across individual patients (Figure S2C), we found a large preponderance of BMDMs over microglia in more aggressive tumor states with increasing proportions of BMDMs correlating with increasing glioma grade (Figure 2A). Strikingly, we found that only two of the 14 myeloid populations were unique to grade IV GBMs while virtually absent in grade 2 and 3 brain tumors (Figure 2B). One was a population resembling early-stage MDSCs (E-MDSCs) that lacked expression of lineage markers, and the other was a population of monocytic MDSCs (M-MDSCs) with stronger expression of CD14. A third myeloid population, designated MAC2, with transcriptomic features of both M1 and M2 macrophages, was also highly enriched in grade IV GBM, though small numbers were observed in Grade 2 and 3 tumors. Transcriptomic analysis of these three populations revealed that E-MDSCs preferentially expressed multiple genes encoding metabolic enzymes, stress-induced genes, and metallothionienes; M-MDSCs exhibited preferential expression of multiple S100A family genes and cellular migration genes; and the MAC2 macrophage population expressed high levels of MHC2, scavenger receptors, and tissue damage associated genes (Figure 2C). No other myeloid population was selectively expressed in GBM.

**Fig. 2.**
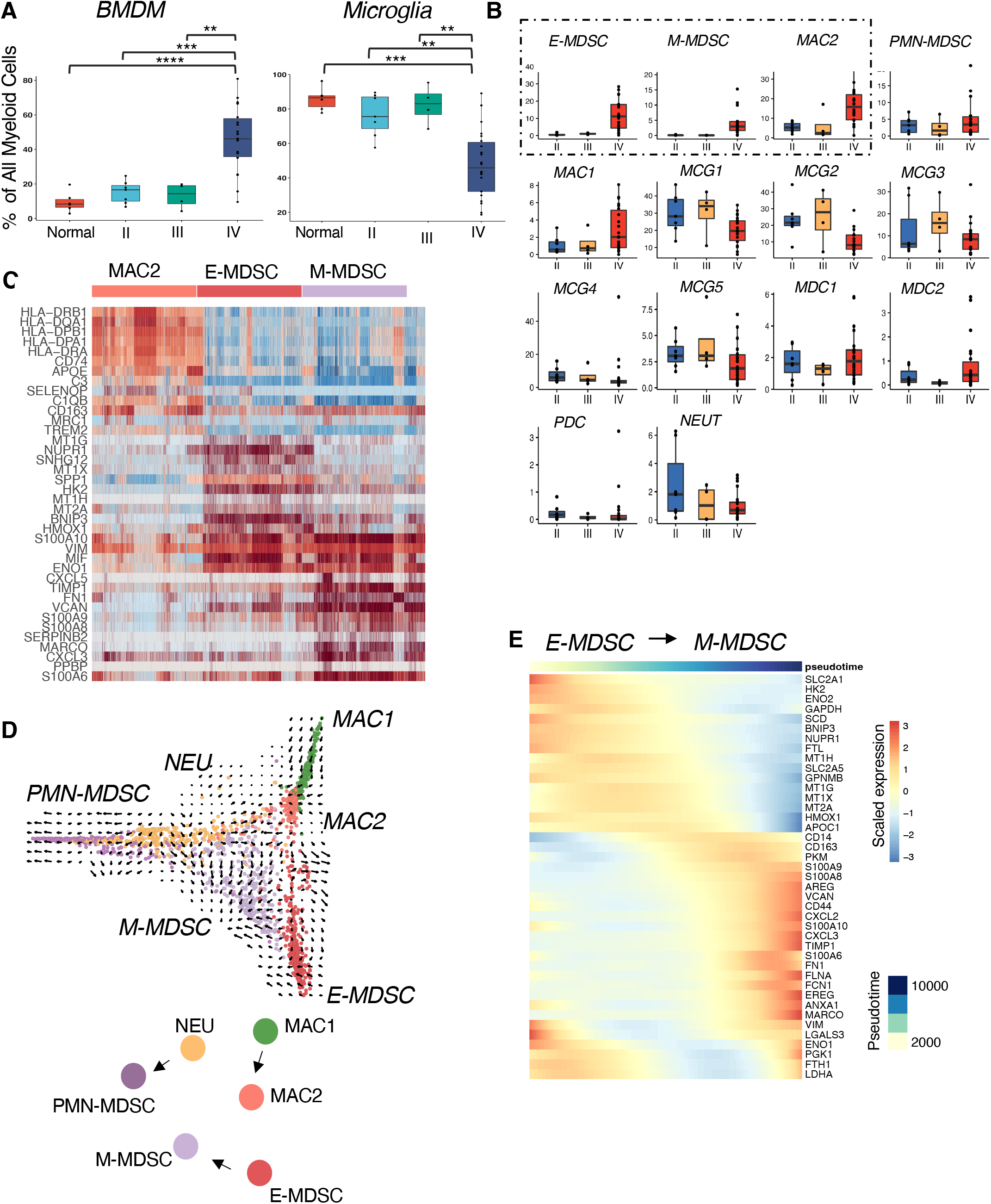
GBM unique myeloid derived suppressor cell exhibit distinct metabolic and functional pathways. (A) The proportion (%) of total myeloid derived cells made up by bone marrow derived myeloid cells and. microglia compared between tumor grades and non-neoplastic tissue. Pairwise comparison and *p* values were obtained using Wilcoxon test. Each dot represents a patient and all data points are shown. Individual data points are superimposed over a Box and Whiskers plot summarizing the data. *p≤0.05, **p≤0.01 ***p≤0.001,**** p≤0.0001 (B) Box plots showing increased proportion of E-MDSC, M-MDSC, and MAC-2 cell populations in grade IV gliomas compared to grade II and III gliomas. Anova test. (C) Heatmap of differentially expressed genes between cells belonging to the three BMDM subsets enriched in glioblastoma demonstrating distinct transcriptomic profiles with each column representing a cell. (D) Diffusion plot with RNA velocity for bone marrow derived myeloid cell clusters in GBM. Cells were randomly downsampled to 300 cells each cluster for visualization. (E) Heatmap demonstrating the pattern of gene expression along the E-MDSC to M-MDSC trajectory.

Although the role of MDSC in inhibition of natural anti-tumor T cell responses in T cell-poor tumors is unclear, none-the-less, a standard test of MDSC function is inhibition of T cell proliferative responses *in vitro*. The GBM-infiltrating MDSC were FACs sorted into M-MDSC (CD14+), PMN-MDSC (CD15+), and E-MDSC (CD14-, CD15-, CD16-) (Figure S3A, Table S3). Cell Trace Violet (CTV) labeled, anti-CD3/CD28 stimulated PBMC were co-cultured independently with each tumor associated MDSC subset at varying MDSC:PBMC ratios. We found that all three MDSC subsets robustly suppress T cell proliferation (assayed by CTV dilution) with M-MDSCs demonstrating the strongest in vitro suppression of both CD4 and CD8 T cell proliferation (Figure S3B-D). These results validate that the MDSC defined transcriptionally in our single cell analysis exhibit a classic functional feature of MDSC.

### E-MDSC, M-MDSC, and TAM exist on a continuum of cellular states

To reconstruct the differentiation trajectory and transcriptional states of the GAMs, we generated diffusion maps and performed pseudotime and RNA velocity analysis on the BMDM cells, specifically focusing on the MDSC and macrophage populations. We found developmental linkage among three pairs of GBM-specific myeloid populations (E-MDSC -> M-MDSC, MAC1 -> MAC2, PMN -> PMN-MDSC). E-MDSCs are the earliest in developmental trajectory and have the potential to develop along a distinct developmental trajectory to M-MDSC (Figure 2D).

Next, in order to identify specific genes that are dynamically changing along the single-cell trajectory from E-MDSC to M-MDSCs, we used pseudotime analysis to evaluate global transcriptomics along individual differentiation trajectories vectorially established through our RNA-velocity analysis. We found that as cells transition between E-MDSC and M-MDSC states, there is increased expression of genes associated with extracellular matrix components and remodeling (*CD44, FLNA, VCAN, FN1*), immune inflammation (*FCN1, S100* proteins), chemokines (*CXCL2, CXCL3*), and monocytic scavenger receptors (*CD14, MARCO, CD163*) generally associated with M2 macrophages. Conversely, there is decrease in expression of genes associated with metabolic pathways including glycolysis (*HK2, ENO2, SCD*), anti-oxidation (*HMOX1, MT1G, MT1H*) pathways, and cellular stress response (*BNIP3, NUPR1*) (Figure 2E, Figure S4). The downregulation of metabolic and hypoxia pathways in the setting of cellular transition from the E-MDSC to M-MDSC states suggests differences in the microenvironmental milieu of the TME that these cells occupy.

### GBM MDSCs exhibit robust catabolic and anabolic metabolism

Given the significant shifts in pseudotemporal expression of genes associated with metabolic pathways as cells transition between different myeloid cellular states, we performed gene set enrichment analysis (GSEA) to further characterize pathways representative of functional states of the MDSC populations. GSEA analysis of differentially expressed genes in E-MDSCs and M-MDSCs compared to other BMDM subsets revealed that the most prominently induced gene sets were dominated by multiple pathways of catabolic and anabolic metabolism including enrichment of glycolysis, oxidative phosphorylation, fatty acid, and mTOR pathways (Figure 3A). The most enriched gene set pathways found in both populations of MDSCs were carbohydrate and lipid metabolism pathways. In addition to glycolysis and fatty acid metabolism, E-MDSCs also exhibited upregulation of other carbohydrate metabolism including fructose and mannose metabolism, the pentose phosphate pathway, nucleotide and amino sugar metabolism, as well as amino acid metabolism. Using SCENIC (*7*), a program that determines the activation state of specific transcription factors based on expression levels of their regulon, we identified E-MDSC-specific activation of transcription factors that regulate metabolic pathways including the *CREB, ATF, PPAR and RXR* family of transcriptions factors critical to homeostasis of glucose, lipid, and amino acid metabolism (Figure 3B). Examining programs more closely in E-MDSCs vs M-MDSCs, E-MDSCs appeared to have higher activation of these transcription factors (Figure 3B), and demonstrated significant increase in expression of genes encoding key enzymes involved in glycolysis: hexokinase-2 (*HK2*) and glyceraldehyde 3-phosphate dehydrogenase (*GAPDH*). This population of cells also showed increased expression of glucose transporter 1 (*SLC2A1*) that allows the influx of glucose into the cell as substrate for glycolysis (Figure S5A-B). These broad anabolic and catabolic transcriptional programs are highly indicative of an active proliferating population. Indeed, E-MDSC and M-MDSC represented the major components of a proliferating myeloid cluster defined by genes such as MKI67. Sub-clustering of cycling myeloid cells demonstrate that E-MDSCs and M-MDSCs represent close to half of the cycling cells (Figure 3C). To validate the major metabolic programs of GBM-associated MDSCs at the protein level, we performed 21 color multi-parametric flow cytometry on ten additional GBM samples (Table S4). Using a combination of antibodies to molecules associated with various myeloid lineages and metabolic enzymes and markers (*8*), we identified two distinct populations of HLA-DR^-^CD33^+^ cells that recapitulated the MDSC subsets identified from the scRNA-seq analysis (Figure 3D). In particular, we could discern an MDSC population that recapitulated high expression of HK2 and GLUT1 similar to the E-MDSC population identified through scRNA-seq and a separate population of MDSCs with higher expression of CD14, CD206, and lower expression of GLUT1 similar to the M-MDSC population identified through scRNA-seq. While both MDSC populations demonstrated upregulation of glucose metabolism, E-MDSCs exhibited higher expression of GLUT1 compared to M-MDSCs. We hypothesized that M-MDSCs may utilize a different glucose transporter. Indeed, gene expression analysis found that they displayed higher expression of GLUT3 transporter, while E-MDSCs displayed higher expression of GLUT1 and GLUT5 transporters (Figure S5C). Our flow cytometry results also demonstrated increased expression of voltage-dependent anion-selective channel 1 (*VDAC1*), mitochondrial import receptor subunit TOM20 homolog (*Tomm20*), and phosphorylated S6 ribosomal protein (pS6) in these cells which are a part of the oxidative phosphorylation and mTOR pathways, respectively, compared to HLA-DR+ macrophages. Taken together, these proteomic studies validated our single cell transcriptomic profiling, demonstrating significant upregulation of multiple metabolic pathways in the MDSCs (Figure S5D). Flow cytometric analysis on matched peripheral blood mononuclear cells (PMBCs) from GBM patients showed that HK2^+^ MDSCs were exclusively found amongst tumor infiltrating myeloid cells and not in the peripheral blood further supporting our hypothesis that these cells alter their metabolic pathways in response to the nutrient-poor TME (Figure 3E. Taken together, these findings demonstrate that GBM MDSCs are highly programed to take up and metabolize nutrients that are limited in the TME for high energy needs as well as rapid cellular expansion via anabolic pathways. These upregulated metabolic programs are characteristic of tumor cells themselves but have not been previously recognized in MDSC in the TME.

**Fig. 3.**
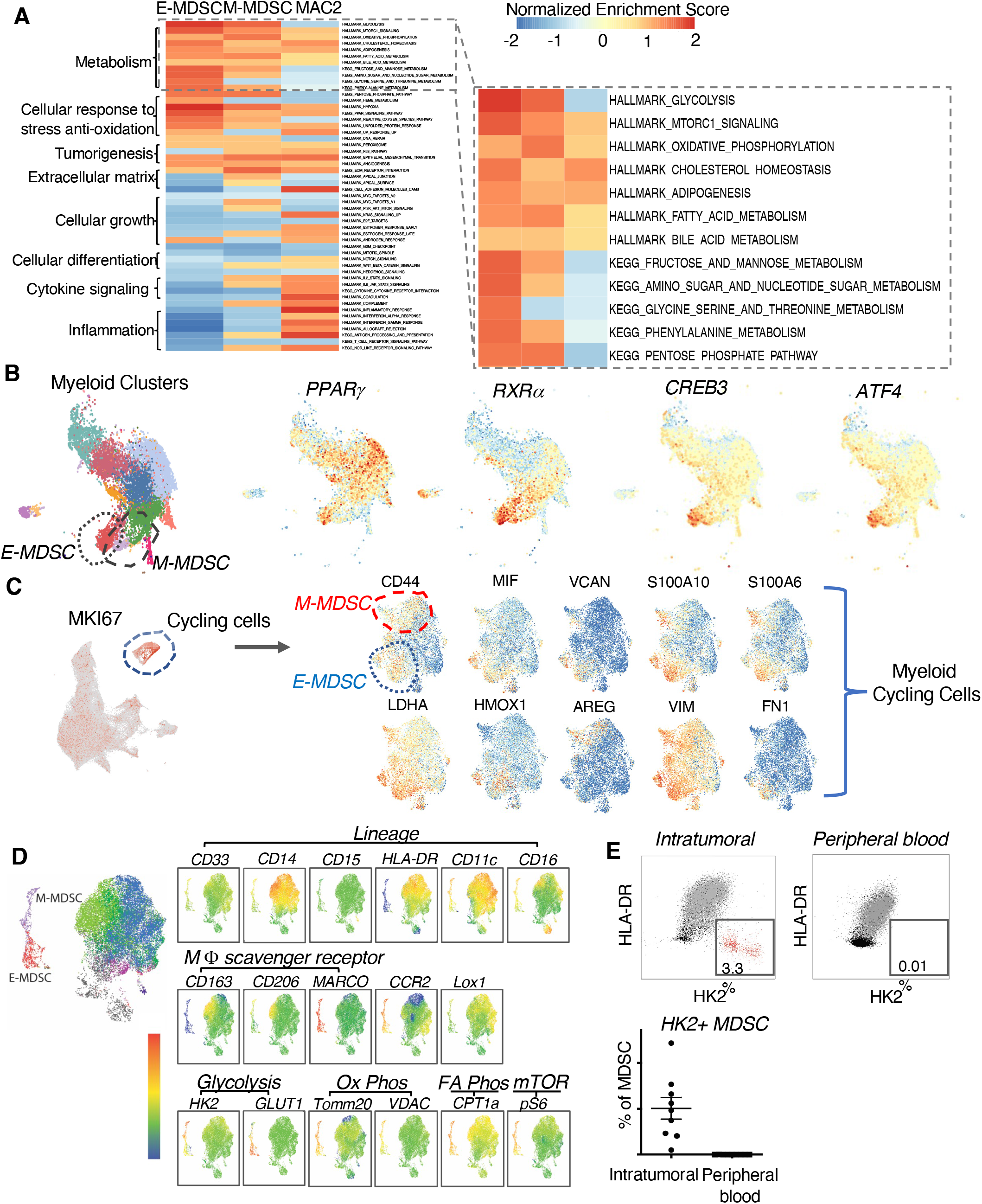
GBM unique myeloid derived suppressor cell exhibit distinct metabolic pathways. (A) Heatmap of Hallmark and KEGG GSEA enriched pathways among E-MDSC, M-MDSC, and MAC2. Enlarged panel demonstrating top upregulated metabolic pathways in MDSCs. (B) UMAP plots demonstrating increased activation of transcription factors that regulate carbohydrate, fatty acid, and amino acid metabolism. (C) UMAP plot demonsirating population of myeloid cells with high expression of cycling gene (MKI67) (left). Subclustering of cycling myeloid cells demonstrate that E-MDSCs and M-MDSCs represent close to half of the cycling cells. UMAP plots of subclustered cycling cells demonstrates large number of cells exhibiting high expression of genes specifically found in E-MDSC and M-MDSC cells (right). (D) t-SNE plots of multicolor flow cytometry illustrating the presence of two populations of myeloid cells with low expression of HLD-DR, high expression of proteins involved in glycolysis, oxidative phosphorylation, and mTOR metabolic pathways. (E) Flow cytometry depicting the unique presence of HK2 positive MDSCs with high expression of signature proteins in metabolic pathways in the tumor microenvironment compared to peripheral blood mononuclear cells.

### E-MDSC co-localize with glioma stem-like cells to pseudopalisading regions of GBM

In order to define potential interplay between the myeloid cells and tumor cells, we next carried out unsupervised clustering analysis of the CD45-cell population (Figure S6A), classifying cells into malignant and non-malignant cell types by inferring copy number variations on the basis of the expression intensity of genes across positions of the genome (inferCNV) (*9*) (Figure S6B). Ten clusters of tumor cells were defined in our UMAP analysis with meta-programs consistent with previous reports (*10*) (Figure 4A). We next evaluated the presence of tumor cell populations across glioma grades and found that tumor populations T3 and T4 are highly selectively expressed in GBMs while virtually absent in grade II or grade III tumors (Figure 4B). To determine potential interactions between myeloid and tumor cells in the GBM TME, we performed pairwise spearman correlation analysis on the proportion of the various transcriptomically determined subpopulations of tumor and immune cells across our patient samples. Strikingly, frequencies of E-MDSC cells were positively correlated in a highly statistically significant way with the presence of only one of the tumor clusters - T4 - across all samples and within GBMs (Figure 4C). Transcriptomic analysis of the T4 tumor populations revealed increased expression of genes associated with angiogenesis (*VEGFA*), neuronal development (*MALAT1*), hypoxia response (*ERO1A*) and glycolysis (*PGK1*). Correspondingly, on GSEA analysis, the T4 tumor cluster demonstrated upregulation of tumorigenesis pathways including VEGF signaling, integrin, glycolysis, and hypoxia pathways (Figure S6C, S6D).

**Fig. 4.**
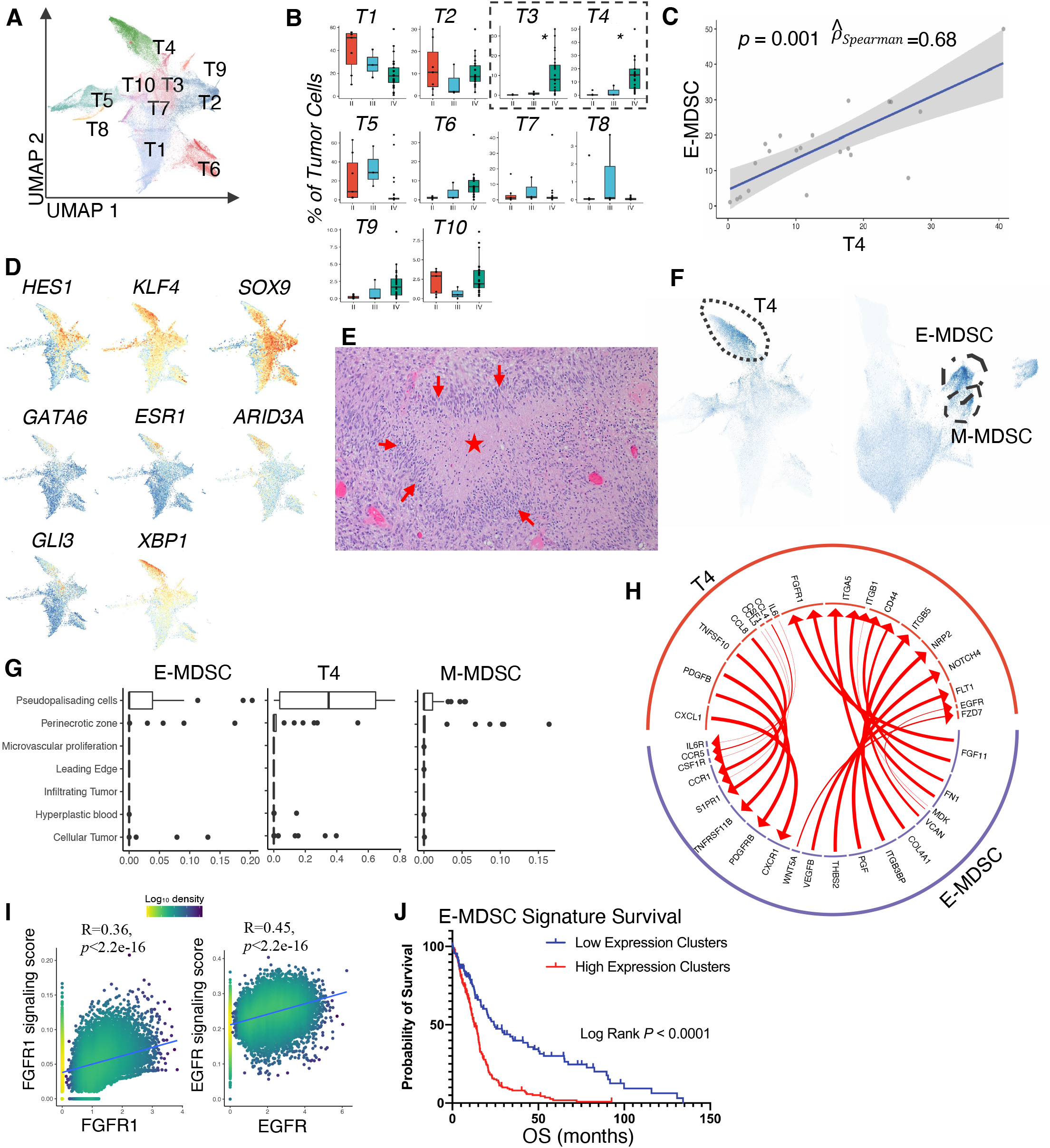
E-MDSCs co-localize with GBM-specific glioma stem cells to pseudopalisading region of tumor. (A) UMAP projection of the expression profile of neoplastic cells demonstrating 10 clusters of tumor cells each delineated by color code. (B) Box plots showing significant over representation of tumor cluster T3 and T4 in GBM comparedto grade III and grade II gliomas. *p≤0.05, **p≤0.01 ***p≤0.001,**** p≤0.0001. Anova. (C) Scatter plot highlighting the spearman correlation between the frequencies of E-MDSC andT4 across GBM patients. (D) UMAP plots demonstrating activation scores of transcription factors that promote cancer stem-ness activated preferentially in glioma stem-cell tumor cluster T4. (E) Hematoxylin and eosin pathological slide of GBM demonstrating the pseudopalisading region characterized by hypercellular cells (arrows) surrounding a central area of necrosis(*). (F) UMAP plot emphasizing the representationof the pseudopalisading gene signature by glioma stem cell cluster T4 and E-MDSCs. (G) Bar plots demonstrating deconvolution of bulk RNA sequencing revealing the representation of T4, E-MDSCs, and M-MDSCs in the pseudopalisading region of GBM. (H) Predicted ligand receptor interactions between MDSC and tumor cell populations in GBMs. (I) Correlation analysis of growth factor expression with gene set signaling score inT4tumor cells demonstrate strong correlation between the expression level of MDSC stimulated growth factor receptor to downstream signaling pathway (J) Kaplan Meier curve illustrating that the E-MDSC expression signature stratified GBM patient survival.

Regulon-based transcription factor activity analysis demonstrated that this T4 tumor cell population upregulated activity of transcription factors previously described in glioma stem-like cells (GSCs) that play key roles in reprogramming differentiated GBM into stem-like cells capable of self-renewal and tumor propagation including HES1, SOX9, and KLF4 (*11*). The T4 population also exhibited activation of ARID3A, GLI3, GATA6, and XBP1 programs, which are less described in gliomas but have been implicated in inducing stem promoting pathways such as Hedgehog signaling pathway and maintaining the stemness of cancer stem cells in other systemic cancers (*12–17*) (Figure 4D). While it is unclear whether GSCs are the one renewable tumor cell giving rise unidirectionally to other tumor subsets, they are a key subset of glioma tumor cells that are responsible for tumor progression, invasion, and resistance to chemoradiation (*18*). Our RNA-velocity analysis of the tumor cell clusters suggests that tumor cluster T4 has the potential to transition to cluster T3 which has the potential to transition into other tumor clusters indicating a stem-like role of the T4 cluster (Figure S6E).

Given that the MDSC and the GSC tumor cell subpopulation shared upregulation of metabolic pathways associated with high energy need in environments of limiting oxygen and nutrient availability, i.e. glycolysis, angiogenesis, and hypoxia pathways, we hypothesized that these cells may geographically colocalize in regions of the tumor requiring upregulation of these pathways to support cellular growth and energy consumption in an environment of limited nutrients and oxygen. We first evaluated the expression level of key genes upregulated among the GBM MDSC subsets, including *HK2* and *HMOX1*, from previously published RNA-sequencing data sets across five major anatomic structures of GBM collected by laser microdissection of tumor tissue (*19*). We found that the expression of *HK2* and *HMOX1* were significantly higher in the pseudopalisading region of GBM compared to other pathologically characterized regions of GBM, including the leading edge, cellular tumor, infiltrating tumor, and microvascular proliferative regions (Figure S7A). The pseudopalisading region of GBM is a hypercellular zone surrounding a necrotic focus (Figure 4E) that is a unique and distinguishing pathological feature of GBM compared to lower grade gliomas. This region is characterized by a hypoxic microenvironment and microvascular proliferation. The presence of pseudopalisading necrosis and microvascular hyperplasia are significant predictors of poor prognosis in gliomas and the hypoxic environment are thought to drive tumor cell survival and stemness (*20*). While these regions have traditionally been thought to be predominately made up of tumor cells, our results suggest that there is significant contribution to the cellular architecture of the pseudopalisading region by the MDSCs.

To determine the contribution of the expression of these genes from MDSCs versus tumor cells, we evaluated the expression of *HK2 and HMOX1* across all three cell populations (tumor, myeloid, and lymphoid cells) and found that the expression of these genes was specifically upregulated in the myeloid population (Figure S7B). To further expand our analysis of the expression of specific marker genes within the pseudopalisading region, we generated a gene signature score derived from genes that are upregulated and downregulated in the pseudopalisading region compared to other anatomical regions in GBM. We discovered that MDSC and T4 tumor cells exhibited the highest pseudopalisading gene signature score (Figure 4F). We further validated the co-localization of MDSC with the T4 tumor subset in the pseudopalisading region by using our scRNA-seq gene expression data to characterize the cell type proportion from bulk RNA-sequencing data across the distinct GBM regions based on morphologic features. We found high composition of tumor cluster T4, E-MDSCs, and to a lesser extent, M-MDSCs in the pseudopalisading region (Figure 4G).

### MDSCs and glioma stem-like cells demonstrate selective upregulation of hypoxia and cellular stress responses

Given the co-localization of the MDSCs and glioma stem cell like tumor cluster T4 to the pseudopalisading region of GBM, we hypothesized that these cells, in addition to their highly metabolic genetic programs, would also exhibit upregulation of functional programs that allow them to persist in this hypoxic niche of the tumor microenvironment. As such, our results further demonstrated that both populations of MDSCs displayed upregulation of genes and transcriptions factors associated with cellular stress response and anti-oxidative response, including hypoxia, reactive oxygen species, and heme metabolism pathways; all of which have previously been implicated to play important roles in in adaptive and innate immune dysfunction and tumorigenesis (*21*) (Figure 5A-B). One particular gene that demonstrated significant upregulation in the E-MDSCs is *HMOX1*, the rate limiting enzyme in the catabolism of free heme and plays a key role in regulating anti-inflammatory and anti-oxidation pathways. Coincidentally, these cells also exhibited increased expression of ferritin light and heavy chains (*FTL, FTH1*) (Figure S5A). On transcription factor analysis, E-MDSCs demonstrated increased activation of transcription factors that regulate the expression of *HMOX1*. In addition to *HMOX1*, E-MDSCs also exhibited increased expression of genes (*NUPR1, ERO1A*) and associated transcription factors linked to the cellular response to hypoxia (Figure 5B). M-MDSCs also upregulated gene pathways involved in hypoxia and stress response as well as upregulation of genes encoding growth factors (*EREG*), chemokines (*CXCL2, CXCL3*) and extracellular matrix (ECM) components that are key facilitators of cell motility.

**Figure 5:**
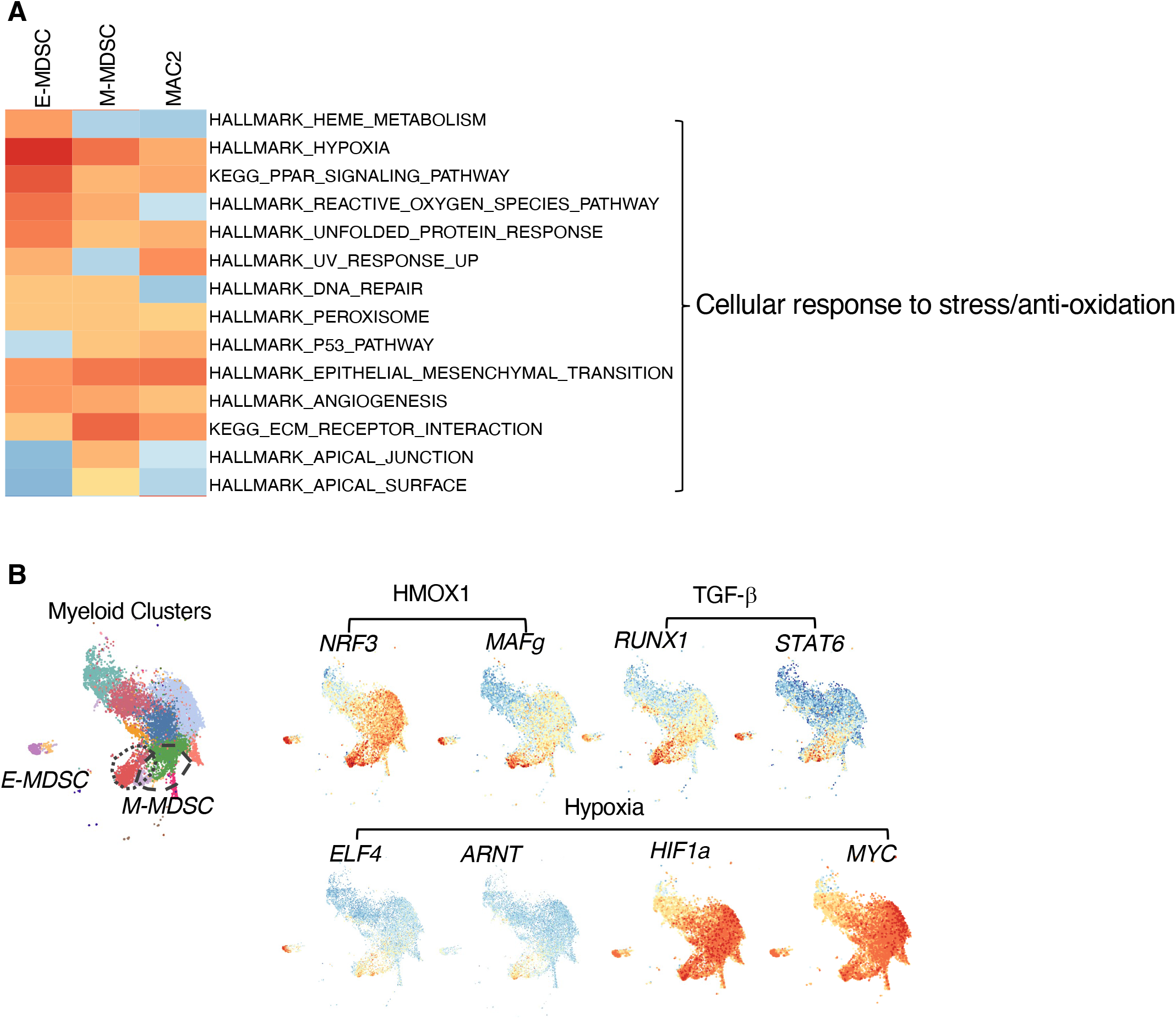
M-MDSC and E-MDSC demonstrate upregulation of hypoxia and cellular stress response pathways. **(A)** Heatmap of Hallmark and KEGG GSEA enriched pathways among E-MDSC, M-MDSC, and MAC2. Enlarged panel demonstrating top upregulated pathways in cellular response to stress and tumorigenesis present in MDSCs. **(B)** UMAP plots demonstrating increased activation of transcription factors that regulate anti-oxidation, hypoxia, and cellular response to stress.

Our results reveal that these distinct populations of E-MDSC and M-MDSC cells uniquely present in GBMs not only upregulate their metabolic programming to competitively survive in a nutrient poor microenvironment, they also upregulate pathways associated with cellular response to low oxygen environment including anti-oxidation and cellular repair pathways to acclimate, thrive and even actively proliferate (Figure 3C) in the hostile milieu of the pseudopalisading region of GBM.

### Cluster interactions indicate a symbiotic relationship between E-MDSC cells and mesenchymal glioma stem-like cells

The co-localization of the E-MDSCs and GSCs to the pseudopalisading region of the GBMs, suggested that short- and mid-range communication, such as from membrane ligands, cytokines, chemokines, growth and differentiation factors, between these cells synergistically drive the aggressiveness and progression of GBMs. We therefore examined cognate ligand-receptor interactions selectively increased between E-MDSC and T4 tumor cells to determine putative modes of communication between these cells. We utilized an established dataset of biologically validated ligand-receptor pairs to score interactions between pairs of cell clusters (*22, 23*) (Table S5). We found remarkably high numbers of ligand-receptor pair interactions between E-MDSC and T4 tumor cells (Figure 4H).

Multiple predicted receptor-ligand interactions suggested an active role for the GSCs in the recruitment and proliferation of E-MDSCs. Our results demonstrated increased chemokine/chemokine receptor signaling from GSCs to E-MDSCs through several ligand-receptor pairs: *CCL8-CCR1, CXCL1-CXCR1,CCL5-CCR1 and CCL4-CCR5*. Some of these interactions have been shown to participate in the recruitment of MDSCs into sites of inflammation and TME in various murine cancer models including gliomas (*24–26*). In addition to these chemokines, IL6-IL6R interactions further enhance MDSC accumulation and induction of genes encoding various immune inhibitory functions (*27*). GSCs also demonstrated increased expression of macrophage colony-stimulating factor 1 (*CSF*), a major myeloid growth factor, paired with high expression of *CSF-1R* on E-MDSCs.

Reciprocally, we identified strong ligand-receptor interactions from E-MDSC to GSCs driving tumor proliferation, survival, and invasion. Extracellular matrix component versican (*VCAN*) is strongly induced on E-MDSCs while its corresponding receptor CD44 is induced in GSCs. E-MDSCs also demonstrated upregulation of ligands for multiple GSC growth factor receptor kinases, including *FGF11-FGFR1*, which promotes tumor cell proliferation and survival through activation of Ras/MAPK, PI3K/AKT, and the phospholipase (PLC) /protein kinase C (PKC) signaling cascade (*28, 29*). Furthermore, E-MDSC demonstrated increased interaction with receptors on T4 tumor cells that are associated with tumor invasion, angiogenesis, and glioma cell migration including integrin a5 (*FN1/ITGA5*) and vascular endothelial growth factor B (*VEGFB/FLT1*) (Figure 4G). Transcription factor analysis of the tumor cells demonstrated increased activation of transcription factors associated with growth factor and angiogenesis pathways including c-Jun, c-Myc and ATF (Figure S8). In order to further validate that growth factors produced by E-MDSCs (Figure 4H) truly are inducing gene programs activated by their cognate growth factors in T4 tumor cells regulated by their cognate receptors, we next generated signaling pathway scores from established gene sets for the key growth factor receptors, EGFR and FGFR1, that demonstrated upregulation in the T4 tumor cell population identified from the ligand-receptor interaction analysis (*30–32*) (Table S6). These signaling scores were then correlated to the expression level of the respective growth factor receptors for each cell in the T4 population. We found strong correlations between the expression level of the growth factor receptor on the cell with the associated signaling score of the growth factor pathway in each cell. This confirms that the inferred ligand-receptor interactions in Figure 4I result in significant upregulation of growth factor pathways driven by MDSCs in T4 tumor cells.

We further evaluated the correlation of the E-MDSC signatures with T4 tumor population signatures amongst TCGA data. We found that the E-MDSC gene signature demonstrated a strong correlation with the tumor T4 population gene signature (Figure S7C). Taken together, the co-localization of E-MDSC and GSC in a specific anatomical site in GBM, together with ligand-receptor interaction mapping support the notion that there is a symbiotic relationship between the E-MDSCs and GCSs in which tumor promotes E-MDSC accumulation and proliferation and E-MDSC accumulation supports tumor growth and invasion.

### MDSC signature is an independent prognostic factor in GBM

To determine whether the presence of these specific populations of MDSCs and GSCs conferred differences in survival amongst GBM patients, we examined the clinical consequences of expression of MDSC and GSC associated genes in bulk TCGA gene expression. We clustered TCGA glioma samples (2013 TCGA, 2016 TCGA) into samples expressing high E-MDSC, M-MDSC, and T4 gene expression signatures (Figure S9A). We found that high expressing samples were 100% GBM while low expressing samples consisted of various percentages of different grades of gliomas (Figure S9B), recapitulating our scRNA-seq results that these populations of cells are unique to grade IV GBMs. Survival analysis demonstrated that patients with high E-MDSC gene expression signature had significantly shorter overall survival compared to patients with low E-MDSC gene expression signature independent of IDH mutation and MGMT methylation status (Figure 4J). Similarly, M-MDSC and T4 gene expression signature also stratified GBM patient survival (Figure S9C).

## Discussion

While MDSC can suppress T cell function in the TME when both are present (*33*), they have also been proposed to play other roles in cancer growth. These alternative functions may be particularly relevant in “cold” tumors where there are very few T cells in the first place. Here we used unbiased scRNA-seq analysis covering more than 750,000 immune cells and 350,000 tumor cells to construct an immune-tumor atlas in gliomas. Our transcriptomic, proteomic, and functional profiling identified two highly proliferative populations of MDSC (E-MDSC, M-MDSC) cells, uniquely present in grade IV GBMs compared to lower grade gliomas. These MDSCs strongly upregulate multiple catabolic and anabolic pathways, and stress and hypoxia response pathways. RNA velocity and ligand-receptor interaction analyses revealed a dynamic continuum of cellular states between MDSCs and intricate cross-talk between MDSCs and tumor cells in GBMs. E-MDSCs co-localize with a specific stem-like tumor subset to the morphologically defined pseudopalisading regions of the cancer. This stem-like tumor subset recruits and activates E-MDSC through production of multiple chemokines and activation factors such as IL6. The E-MDSC reciprocally promote stem-like tumor cell growth and mesenchymal differentiation through secretion of growth and angiogenetic factors.

In gliomas, GBM patients were found to have significant elevation of MDSCs in the periphery with increased levels of MDSCs being associated with worse prognosis (*34, 35*). However, the distinct transcriptomic profile of subsets of MDSCs in GBMs and the precise mechanism by which MDSCs traffic to the tumor to promote tumor growth and immunosuppression are not known. We identified three populations of MDSCs in GBMs including E-MDSCs, M-MDSC, and PMN-MDSCs with GBMs being significantly enriched in E-MDSCs and M-MDSCs. Both populations of cells, though particularly E-MDSCs, exhibited robust and diverse set of metabolic genes with upregulation of glycolysis, oxidative phosphorylation, and mTOR pathways, indicating programming for high energy utilization and anabolism necessary for rapid expansion of the population. There has been much focus on genetic and epigenetic alterations in cancer cells themselves that facilitate adaptation of their metabolism to allow proliferation in nutrient scarce microenvironments and out-compete the non-transformed cells in the TME. Under conditions of limited glucose, cells can reprogram their metabolism towards other pathways including oxidative phosphorylation and mTORC1 signaling to drive energy generation (*36, 37*). Under lower oxygenation, tumor cells activate cellular stress responses to mitigate the negative impact of oxygen deprivation and preserve proliferative capacity. Interestingly, these same pathways are also among the highly induced pathways in the MDSC population (in particular E-MDSC) from our GSEA analysis.

In addition to upregulation of additional metabolic pathways, in order to maintain their proliferative expansion and evade apoptosis to thrive in the harsh hypoxic environment of the pseudopalisading region, E-MDSCs also reprogram their transcriptomics to handle hypoxia and oxidative stress, highly expressing metallothionein genes (*MT2A, MT1G*) and genes involved in heme degradation (*HMOX1, FTL, FTH1*). All of these programs are hallmarks of tumor-intrinsic biology but have not been appreciated to be so highly expressed in and functionally relevant for geographically localized MDSC populations. Metallothioneins function as potent antioxidants through scavenging of free radicals and have been associated with increasing glioma grade and worse survival (*38*). *HMOX1* is the rate limiting enzyme in the catabolism of free heme and plays a key role in regulating anti-inflammatory and anti-oxidation pathways. Our results indicate that MDSCs have similar compensatory mechanisms to actively reprogram their metabolism to ensure survival in the hypoxic TME.

While GBM rarely metastasizes systemically, it remains one of the most recalcitrant tumors due to its aggressive invasion and infiltration into surrounding normal brain tissue, making surgical cure impossible, with recurrence inevitable despite treatment. Prior studies have suggested that a hypercellular zone of cells migrating away from a necrotic focus, defined pathologically by the pseudopallisading region pathognomonic of GBM, may be responsible for the highly invasive nature of GBM. While prior studies have shown that the cells in this zone secrete high levels of proangiogenic factors that promote tumor growth (20), the origins of these factors have not been defined. Our scRNA-seq results revealed that the most selectively expressed genes in the pseudopalisading region from prior bulk RNA-sequencing data were contributed by E-MDSC and a specific stem-like tumor cell subset with a high epithelial-mesenchymal transition (EMT) signature. We hypothesize that this hypoxic core leads to tumor cell migration and microvascular proliferation at its periphery resulting in the recruitment of MDSCs from the peripheral blood to the TME. Our data demonstrated that the glioma stem cell populations support the development and trafficking of the MDSCs through *CSF1/CSFR, CCL8/CCR1, and CXCL1/CXCR1* signaling. And reciprocally, the MDSCs support the growth and invasion of the glioma stem cells in the hypoxic environment through engagement of growth factor (*FGF11/FGFR1, VCAN/EGFR*) and cellular migration (*FN1/CD44*) pathways.

The cellular components of the GBM immune microenvironment have often been viewed in a static state of terminal differentiation. Using computational lineage tracing, we discovered that the myeloid cells in the TME represent a dynamic continuum of cellular states. Furthermore, cells in immature states, such as E-MDSCs, have the potential to transition into M-MDSCs. This finding indicates that not only do E-MDSCs have the capacity to promote tumor progression and immune suppression, they also have the capability to transition into other cellular states that also have the ability to promote tumor growth and T cell suppression through distinct but complementary mechanisms. This suggests that targeting one population of MDSCs might be suboptimal in counteracting their effect on GBM progression, while targeting multiple populations are required to effectively address the immunosuppressive GBM TME.

In summary, our large-scale single-cell analysis elucidated unique MDSC populations as key facilitators of glioblastoma progression and mediators of tumor immunosuppression. Our results for the first time characterized the phenotypic and functional heterogeneity of MDSC populations in GBM with each population exhibiting unique gene signatures and phenotypic markers that differentiate them from other bone marrow derived myeloid cells. We further illuminated the molecular mechanisms that govern the recruitment and metabolic programs that allow the accumulation of MDSCs in the GBM TME along with specific ligand-receptor interactions between MDSCs and glioma stem cells that likely contribute to their role in promoting tumor invasion. Our comprehensive analysis will lay the foundation for the generation of novel strategies to target these specific myeloid compartments and their metabolism as promising therapeutic intervention in this deadly cancer.

## Acknowledgments

We thank the members of the Experimental and Computational Genomics Core, supported by NCI Cancer Center Support Grant P30 CA006973

## Funding

National Institutes of Health grant F32 (CJ)

Neurosurgery Research Education Foundation (CJ)

Bloomberg Philanthropies (DP)

## Author contributions

Conceptualization: CJ, ML, DP

Methodology: CJ, CC, ZJ, WH, RL, HZ, ML, HJ, DP

Investigation: CJ, DB, AD, ET, MZ, HZ, JC

Visualization: CJ, CC, AD, ET, ZJ, WH, RL

Funding acquisition: CJ, DP, DE, HJ, JP, CB, ML

Project administration: CJ, JE, HJ, DP

Supervision: ML, CB, JP, JE, HJ, DP

Writing – original draft: CJ, CC, AD, ET, SY, DP

Writing – review & editing: CJ, CC, SB, AD, ET, MZ, ZJ, WH, RL, HZ, JC, FR, JW, SY, ML, CB, JP, JE, HJ, DP

## Competing interests

None

## Data and materials availability

All single-cell RNA-seq data will be deposited at the NCBI’s Gene Expression Omnibus (GEO) and will be publicly available at the date of publication. All other data are available in the main text or the supplementary materials.

## Supplementary Materials

Materials and Methods

Supplementary Text

Figs. S1 to S9

Tables S1 to S6

References (*1–38*)

## Notes

### Competing Interest Statement

The authors have declared no competing interest.

