## Supplemental Tables and Figures for "Distinct Myeloid Derived Suppressor Cell Populations Promote Tumor Aggression in Glioblastoma"

Supplementary Materials for

Single Cell RNA-Sequencing Identifies Distinct Myeloid Derived Suppressor Cell Populations Promoting Tumor Aggression in Glioblastoma

**This PDF file includes:**

Materials and Methods

Figs. S1 to S19

Tables S1 to S6

Materials and Methods

Human subjects

Patients were prospectively enrolled into this study at Johns Hopkins Hospital. Written informed consent was provided by all participants according to approved Institutional Review Board Protocol. Fresh tumor tissue and blood were collected at the time of surgery following confirmation of primary glial neoplasm on frozen section. None of the patients was treated with chemotherapy or radiation prior to tumor resection. Tumor samples span all glioma stages (Grade II, Grade III, and Grade IV). Patients undergoing surgical resection for seizure focus were consented for collection of non-malignant brain tissue as control.

Tumor dissociation

Patient tumors or non-malignant brain tissue were mechanically cut into about 1mm^3^ tissue in DMEM medium (Gibco). They were then enzymatically digested with MACS tumor dissociation kit (Miltenyi Biotec) on the gentleMACS Dissociator (Miltenyi Biotec) per manufacture instructions. Dissociated cells were filtered by a 70-mm strainer and centrifuged at 300g for 7 min. After removing the supernatant, the pelleted cells were suspended in red blood cell lysis buffer (Miltenyi Biotec) for 2 min to remove red blood cells. Tumor samples with significant extent of myelin underwent myelin removal process using magnetic Myelin Removal Beads (Miltenyi Biotec) on the autoMACS (Miltenyi Biotec). The cells were then washed with sorting buffer (PBS supplemented with 2% FBS) and accessed for viability using trypan blue exclusion.

Peripheral blood mononuclear cell collection

Peripheral blood mononuclear cells (PMBCs) were isolated after Ficoll-Paque centrifugation. Briefly 80ml of fresh peripheral blood was collected at the time of surgery in EDTA anticoagulant tubes and layered on top of Ficoll-Paque after twofold dilution with phosphate-buffered saline (PBS). After centrifugation (2200 rpm, 20 minutes, no brake), PBMCs are collecting from the mononuclear cell band layer. PBMCs are viability frozen in 10 million per mL until future use.

Fluorescence-activated cell sorting

Single cell suspensions of the tumor and non-malignant brain tissue were stained with antibodies for 30 minutes at 4°C against CD45+ (PE, BD Bioscience), CD3+ (APC, BD Bioscience) and nuclear stain DyeCycle Violet (Invitrogen) for FACS sorting, performed on Beckman Coulter MoFlo XDP. Cells were sorted into three live populations, T cells (CD45+CD3+), non-T immune cells (CD45+CD3-), and non-immune cells (CD45-, DyeCycle Violet+) directly into cold PBS + 1% bovine serum albumin (BSA, Sigma-Aldrich) with a post-sort purity of >98%. Sorted cells were then counted and assessed for viability using trypan blue and re-suspended in 700cells/ml in PBS + 0.04% BSA.

Single Cell Sequencing

scRNA-sequencing of myeloid and tumor cells were performed using the 10X Single Cell 3’ Gene Expression Kit v3 and scRNA-seq and scTCR-seq of T cells were performed using the 10X Single Cell 5’ Immune Profiling Kit v1.1 (10X Genomics, Pleasanton, CA, USA). Cells were captured in droplets at a targeted cell recovery of 5000-10,000 cells per lane. scRNA library generation was performed per protocol. Following cell barcoding in droplets and reverse transcription, emulsions were broken and cDNA purified using Dynabeads. cDNA was amplified by 11 PCR cycles for 3’ gene expression and 13 PCR cycles for 5’ immune profiling. Amplified cDNA was then used for 3’ gene expression library construction or 5’ gene expression library construction and TCR enrichment. Sequencing was performed using an Illumina NovaSeq 6000 with 310 million reads per sample and a sequencing configuration of 26x8x98 (UMI × Index × Transcript read). The Cell Ranger 3.0.2. pipeline software (10X genomics, California) was used to align reads and generate expression matrices for downstream analysis.

MDSC Suppression Assay

MDSCs were isolated from glioma tumors using fluorescence-activated cell sorting. Tumor associated myeloid cells were harvested through tumor dissociation. Surface markers were directly stained with the following fluorochrome-conjugated antibodies (Table S5). Cell viability was assessed using Propidium Iodide (Stem Cell Technologies). Cells were sorted using MoFlow XDP directly into cold RPMI supplemented with 10% heat-inactivated fetal bovine serum (FBS, ThermoFisher) and 10mg/ml gentamicin (ThermoFisher). Cells were then washed, counted, and resuspended at concentration for suppression assay.

Cell Trace^TM^ Violet-labeled responder peripheral blood mononuclear cells (PBMC) from a healthy donor were cocultured with different effector to target ratios with unlabeled tumor MDSCs sorted from patients with glioma. T cell proliferation was induced by addition of anti-CD3/anti-CD28 coated microbeads (Dynabeads Human T Activator, ThermoFisher) at a bead-to-cell ratio of 1:4. Cells were cultured for 96 hours in 96-Ubottom plate in complete RPMI medium (Gibco). Proliferation was assessed using Cell Trace^TM^ Violet dilution detected by a Celesta flow cytometer. Relative proliferation was calculated against stimulated responder PBMC that were cultured without MDSC. Data is representative of this assay done multiple times with different patients all displaying similar results.

ScRNA-seq filtering and normalization

Lymphoid, myeloid, and tumor data sets were processed separately. After combining raw counts from the filtered output of CellRanger, we further filtered cells based on both minimum number of genes and counts. Selection of each was performed using histograms of each parameter by cell (Figure S1A). In each case, a minimum threshold of 500 genes and 750 UMI counts was sufficient to exclude populations of low-quality cells visible as the lower portion of the bimodal/multimodal distributions visualized in the histogram. Genes expressed in fewer than .1% of the cells were also removed. Finally, counts were normalized to the total UMI count by cell and log-scaled using Seurat.

Data integration

Data integration was performed using Seurat v3 using reciprocal principal component analysis to project cells into a shared space followed by identification of anchors for mutual nearest neighbor integration. One female and one male patient were selected as references for integration.

Scaling, principal component analysis, clustering, and dimensional reduction

Scaling of both corrected and uncorrected normalized gene expression values was calculated, although the uncorrected values were used only for visualization of differential gene expression with heatmaps. In both cases, values were scaled to a mean of zero and a standard deviation of one by gene. The scaled, corrected values were used for principal component analysis, after which 60 principal components were selected based on elbow plot for all data sets. The 60 principal components were used for dimensional reduction by uniform manifold approximation projection (UMAP) as well as generation of a shared-nearest neighbor network followed by Louvain clustering. Clusters with very similar gene expression profiles were combined and in some cases, clusters were isolated for further clustering.

Differential gene expression

Library size normalized gene expression matrix was imputed using SAVER (*75*) to address potential dropouts, and the imputed values are log2-transformed after adding pseudocount of 1. A linear mixed-effect model is used to identify genes that are significantly differential between cell clusters or between two groups of samples within each cell cluster. The details of the mixed-effect model can be viewed at the following GitHub page: https://github.com/zji90/Raisin/. The p-values are adjusted for multiple testing using the Benjamini-Hochberg procedure. Genes with FDR<0.05 are considered to be significant differential genes.

Cell cycle scoring

Seurat’s CellCycleScoring was used to identify cells with high G2M or S phase contributions. While this was not used for regression of gene expression values during scaling of all cells, it was used to identify clusters of cells undergoing mitosis. These clusters were isolated and then scaling was performed on these cells alone with regression of cell cycle scores. Finally, principal component analysis and clustering was performed on the regressed, scaled values to attempt to identify types of cells present in the cycling clusters.

Calculation of transcription factor signaling networks

The SCENIC workflow was first used to identify transcription factor activities by cell. All data sets were pooled and then 50,000 cells sampled from the target metadata due to memory limits. Transcription factor scores were then correlated with receptor expression after exclusion of receptors which were predicted targets of specific transcription factors. These correlations were used to build a network between transcription factor and receptor. Finally, ligand pairs for predicted receptors were identified using CellphoneDB2. After generation of networks for high and low grade tumors, the two networks were compared to identify transcription factors and receptors predicted to be activated in only high or low grade tumors.

RNA velocity analysis

RNA velocity analysis was performed as previously described sing the velocyto.py.python package for annotating transcripts as spliced or unspliced, followed by the velocyto.R R package to perform velocity estimation. Briefly, transcripts are marked as either spliced or unspliced based on the presence or absence of intronic regions in the transcript. For each gene, a simple model of RNA dynamics is then fit to the data. Last, the RNA velocity is estimated for each cell by looking for over- or underrepresentation of spliced to unspliced ratios. RNA velocity is visualized on a diffusion plot, with vector fields representing the averaged velocity of nearby cells. (*76*)

### Pseudotime analysis

We apply TSCAN (v.1.7.0) to reconstruct the pseudotime on diffusion maps space for the cells of six cell types (E-MDSC, M-MDSC, MAC1, MAC2, PMN-MDSC, NEUT) from untreated and tumor-grade-IV patients. Our method identifies genes or transcripts that variate along a pseudotime trajectory with statistical significance. We consider the gene expression pattern along a pseudotime trajectory as functional data and represent the data using B-spline basis. We describe the expression of gene *g* along pseudotime $t=1,\ldots,T$ in a hierarchical model. The first hierarchy is to describe the population-level pattern as $x_{s}^{T}\beta_{ig}$ where $x_{s}$ is the design matrix for sample s, and $\beta_{ig}$ is the coefficient w.r.t the i-th B-spline basis and gene g. To test whether a gene variates along a pseudotime trajectory, we let $x_{s}=I_{1}$ and $\beta_{ig}\in R$. It is straightforward to see that the population pattern on all basis is $X_{s}^{T}\beta_{g}$ where $X_{s}=I_{N}\bigotimes x_{s}$, $\beta_{g}=\left[ \beta_{1g},\beta_{2g},\ldots,\beta_{Ng} \right]^{T}\in R^{N}$, and N is the number of B-spline basis. The second hierarchy is to describe the sample-level pattern as $x{{}_{s}^{T}\beta}_{ig}+u_{igs}:=a_{igs}$ where $u_{gs}\sim N\left( 0,\tau_{g} \right),\forall i$, $a_{igs}\sim N\left( x_{s}^{T}\beta_{ig},\tau_{g} \right)$. The third hierarchy is to describe the observed expression of gene g in cell c of sample s as$e_{gcs}=\sum_{i=1}^{N} \phi_{i}\left( t_{gcs} \right)a_{gs}+\epsilon_{gcs}$ where $\epsilon_{gcs}\sim N\left( o,\sigma_{gs}^{2} \right),\forall c.$

The parameters are $\theta=\left( \tau_{g},\sigma_{gs}^{2},\beta_{g} \right)$where $\tau_{g}\in R^{S\times S}, \sigma_{gs}\in R,\beta_{g}\in R^{N},$S is the number of samples (patients); the latent variable is $u_{gs}\in R^{N}$ and $\varphi_{cs}\in R^{N\times1}$ is the B-splines basis function of N basis.

We apply Expectation-Maximization (EM) algorithm to solve this problem and obtain the estimated $\beta_{g}$ for further statistical testing. We perform permutation test where we permute the pseudotime order of the cells and bootstrap the cells within each sample for 100 times. In each permutation, we rerun the above model and redo EM. The number of knots (range from 0 to 30) are automatically selected using BIC. Maximal EM iteration is 100. Convergence cutoff is 1. The null is that all elements in $\beta_{g}=\left[ \beta_{1g},\beta_{2g},\ldots,\beta_{Ng} \right]^{T}\in R^{N}$ are 0. We consider the *p*-value as the percentage of permutation (out of 1000 times) that have log-likelihood greater than the original (pseudotime is not-permuted) log-likelihood. We apply Benjamini-Hochberg (BH or its alias fdr) method to adjust multiple testing. We consider genes with adjusted *p*-values < 0.05 as trajectory differential with statistical significance. We apply *k*-means clustering to group trajectory differential genes with similar pseudotime pattern using standardized gene expression levels. The averaged model-fitted values (fitted resolution = 1000) on standardized scale of the genes within each cluster are used to represent the cluster pattern.

Immune-metabolic ex vivo flow cytometry staining

All flow cytometry antibodies used for phenotypic and metabolic analysis can be found in Table S3. Cryopreserved PBMCs or tumor single cell suspensions were thawed in RPMI (Gibco) + 20% FBS (Atlanta Biologicals). Cells were washed once in PBS and immediately stained for viability with Biolegend Live/Dead Zombie NIR fixable Viability Dye and BD Fc Block^TM^ for 10 min at room temperature. Cell surface staining was performed in 100ul of 20% BD Horizon^TM^ Brilliant Stain Buffer + PBS with surface stain antibody cocktail for 20 minutes at room temperature. Cells were washed with 1X Permeabilization/Wash buffer. Intracellular staining (ICs) was performed in 100ul of 1X Permeabilization/Wash buffer with ICS antibody cocktail for 45 minutes at room temperature. Cells were washed once with Permeabilization/Wash buffer, then resuspended in 1% Paraformaldehyde for acquisition by flow. Samples were run on a 3 laser Cytek Aurora spectral flow cytometer. FCS files were analyzed using Flowjo v10 (10.6.2.) software. High-dimensional unbiased analysis of cell phenotypes was performed using FlowJoDownsample v3 and UMAP.

Cell-cell interaction analysis/ Ligand-receptor interaction analysis

*Pseudo-bulk gene expression data.* We first obtained the pseudo bulk gene expression matrix for each patient as follows: aggregate the raw read count within each cell cluster, normalized the data by count per million (CPM), then log2 transformed the data after adding pseudo count 1. Pseudo-bulk gene expression matrices are generated with each matrix corresponding to a patient sample. Within each pseudo-bulk matrix, each row represents a gene and each column represents a cell type.

*Normalized interaction score.* We utilized a curated list of immune related and non-immune related ligand-receptor pairs from previously published literature (Table S4). The normalized interaction score for a specific ligand-receptor pair between cell type and cell type B is defined as the product of min-max normalized pseudo-bulk expression levels of ligand in cell type A and that of receptor in cell type B.

$$NIS\left( \begin{aligned} ligand,cell type A, \\ receptor, cell type B \end{aligned} \right)=norm\_pb\_expr\left( ligand, cell type A \right)\times norm\_pb\_expr\left( receptor, cell type B \right)$$

$NIS$ = normalized interaction scores

$Norm\_pb\_expr(ligand, cell type A)$ = min-max normalized pseudo-bulk expression levels of ligand in cell type A

$Norm\_pb\_expr(receptor, cell type B)$ = min-max normalized pseudo-bulk expression levels of receptor in cell type B

Specifically, for each patient sample, we first divided the pseudo-bulk gene expression matrix into three subsets based on the cell population, i.e. myeloid, tumor, and T cells. We then performed the min-max normalization across each gene within each cell population. Via this normalization, for each gene, we assigned a value of 1 to the cell type with the highest expression level and a value of 0 to the cell type with the lowest expression level. This normalization method allowed us to identify ligand and receptor pairs that were highly expressed in specific cell types within the larger cell population, rather than those genes that were universally highly express across all cell types.

$$f\left( x_{gijk} \right)= \frac{x_{gijk}-\min_{1\leq l\leq K_{ij}} (x_{gijl})}{\max_{1\leq l\leq K_{ij}} (x_{gijl})-\min_{1\leq l\leq K_{ij}} (x_{gijl})}$$

Here $f$is the min-max normalization function, $x_{gijk}$ is the pseudo bulk expression levels of gene $g$, patient sample $i$, cell population $j$, cell type $k$. $K_{ij}$ is the number of cell types within patient sample $i$ and cell population $j$, and min and max operations are iterated over all possible cell types $1\leq l\leq K_{ij}$

*Identifying significant cell clusters and ligand-receptors.* We searched through all possible cell cluster pairs, i.e. pairwise combinations between 16 T cell clusters, 10 tumor cell clusters and 18 myeloid cell clusters. For each cluster pair, we summarized the number of significant ligand-receptor pairs whose interaction score could differentiate tumor grade IV patients from lower grade (grade II, III). For each ligand-receptor pair, Wilcoxon rank-sum test was performed to compare the normalized interaction scores between grade IV patients and lower grade patients. We also defined the enrichment score as the difference between average normalized interaction scores in grade IV patients and lower grade patients. Benjamin- Hochberg method was applied to adjust for multiple testing, and ligand-receptor pairs with FDR<0.05 and enrichment score >0.3 were reported as significant.

*Ligand-receptor cross-sample correlation analysis.* For each cell cluster pairs of interest, we also detected ligand-receptor pairs whose expressions were highly correlated across grade IV GBM patients. Spearman correlation was calculated and circle plots were made using the *circlize* package (*77*).

*TCGA correlation analysis.* The TCGA Glioblastoma (GBM) gene expression RNA-seq count data were downloaded from UCSC Xena, which included 173 samples. For each sample, we defined the gene signature scores for a cell cluster by the average expression values of top differentially expressed genes (FDR< 0.05 & log_2_ fold change <0.5. We then calculated the Pearson correlation between signature scores of myeloid cell clusters and T cell clusters, as well as myeloid cell clusters and tumor cell clusters. To account for the  potential effect of CD45+ cell levels, we also calculated the adjusted gene signature scores, i.e. dividing the original signature scores by the expression levels of *PTPRC* (CD45).

*Correlation of receptor level with induced gene set* programs. Genes associated with signaling of cognate receptors identified from ligand-receptor interaction analysis were identified using established gene sets from Molecular Signatures Database. These were used to generate cell level scores with UCell [1]. The resulting scores were linearly correlated (Pearson correlation) with normalized RNA expression for the paired receptor and visualized with a pseudocolor plot.

Deconvolution of Ivy bulk RNA-sequencing using scRNA sequencing

BAM files of bulk RNA-seq data from different pathological regions of glioblastoma were obtained from the Ivy Glioblastoma Atlas Project (https://glioblastoma.alleninstitute.org/) and gene expression read counts are obtained using StringTie (v1.3.6) (*78*). MuSiC was performed to deconvolute cell type proportions of bulk RNA-seq samples using the clusters identified in the single-cell RNA-seq data (*79*).

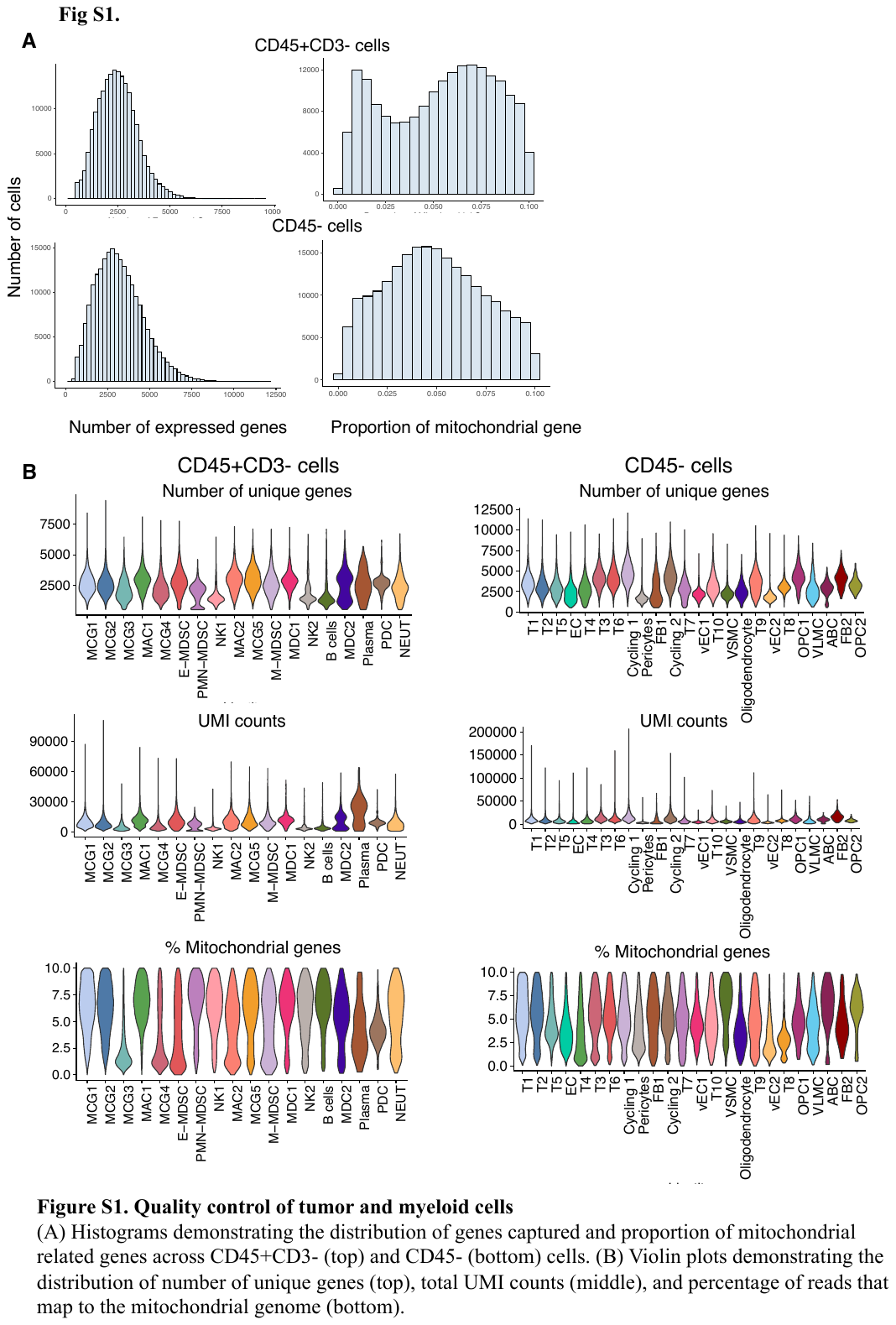

**Fig. S1. Quality control of tumor and myeloid cells**

(A) Histograms demonstrating the distribution of genes captured and proportion of mitochondrial related genes across CD45+CD3- (top) and CD45- (bottom) cells. (B) Violin plots demonstrating the distribution of number of unique genes (top), total UMI counts (middle), and percentage of reads that map to the mitochondrial genome (bottom).

**Fig. S2. Single cell atlas of CD45+CD3- cells in gliomas**

(A) UMAP plot of CD45+CD3- cells in gliomas demonstrating 14 clusters of myeloid lineage cells, two clusters of natural killer cells, one cluster of B cells, and one cluster of plasma cells. (B) Violin plots of expression level of canonical microglia genes across myeloid cell clusters. (C) Stacked bar plots highlighting the variation in proportion of immune cell populations among patients.

Figure S3. E-MDSCs and M-MDSCs demonstrate T cell suppression

A) Multicolor flowcytometry gating strategy to isolate MDSC subsets E-MDSC (HLA-DR-CD33+CD14-CD15-CD16-), M-MDSC (HLA-DR-CD33+CD14+), PMN-MDSC (HLA-DR-CD33+CD15+LOX1+) from fresh single cell suspension of GBM tumor samples. (B) M-MDSC (left) and E-MDSC (right) were plated at a 1:1 ratio with healthy donor PBMC demonstrating suppression (red) when compared to PBMC cultured without MDSC (blue) as determined by differing cell trace violet (CTV) dilution. (C) M-MDSC suppression of CD4 and CD8 T cells quantified by percent divided and percent suppressed at varying M-MDSC:PBMC ratios. (D) M-MDSC suppression of PBMC demonstrated by M-MDSC:PBMC ratio titration from 1:1 (left), 1:4 (middle), and 0:1 (right) and CTV dilution.

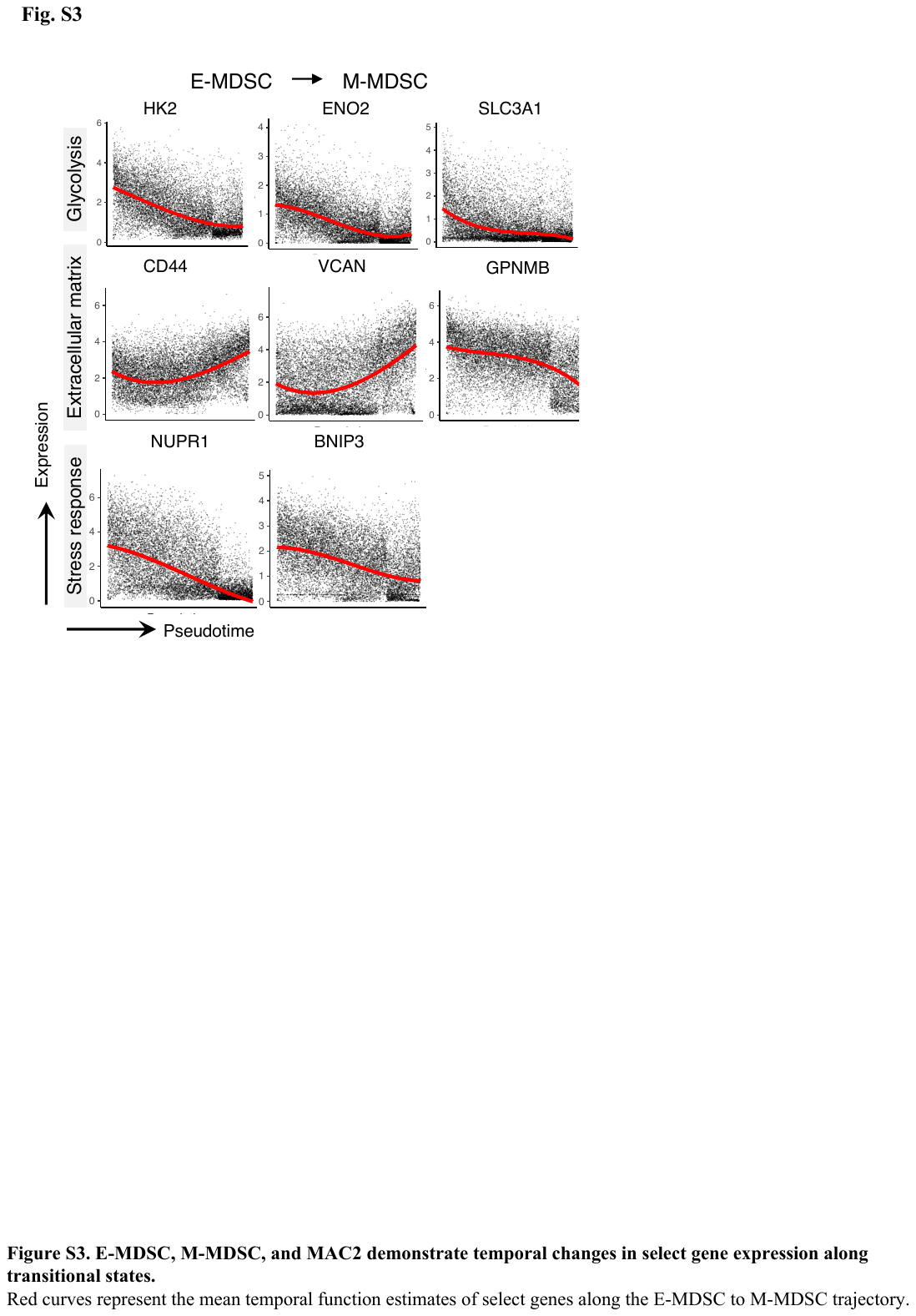

Fig. S4. E-MDSC, M-MDSC, and MAC2 demonstrate temporal changes in select gene expression along transitional states.

Red curves represent the mean temporal function estimates of select genes along the E-MDSC to M-MDSC trajectory.

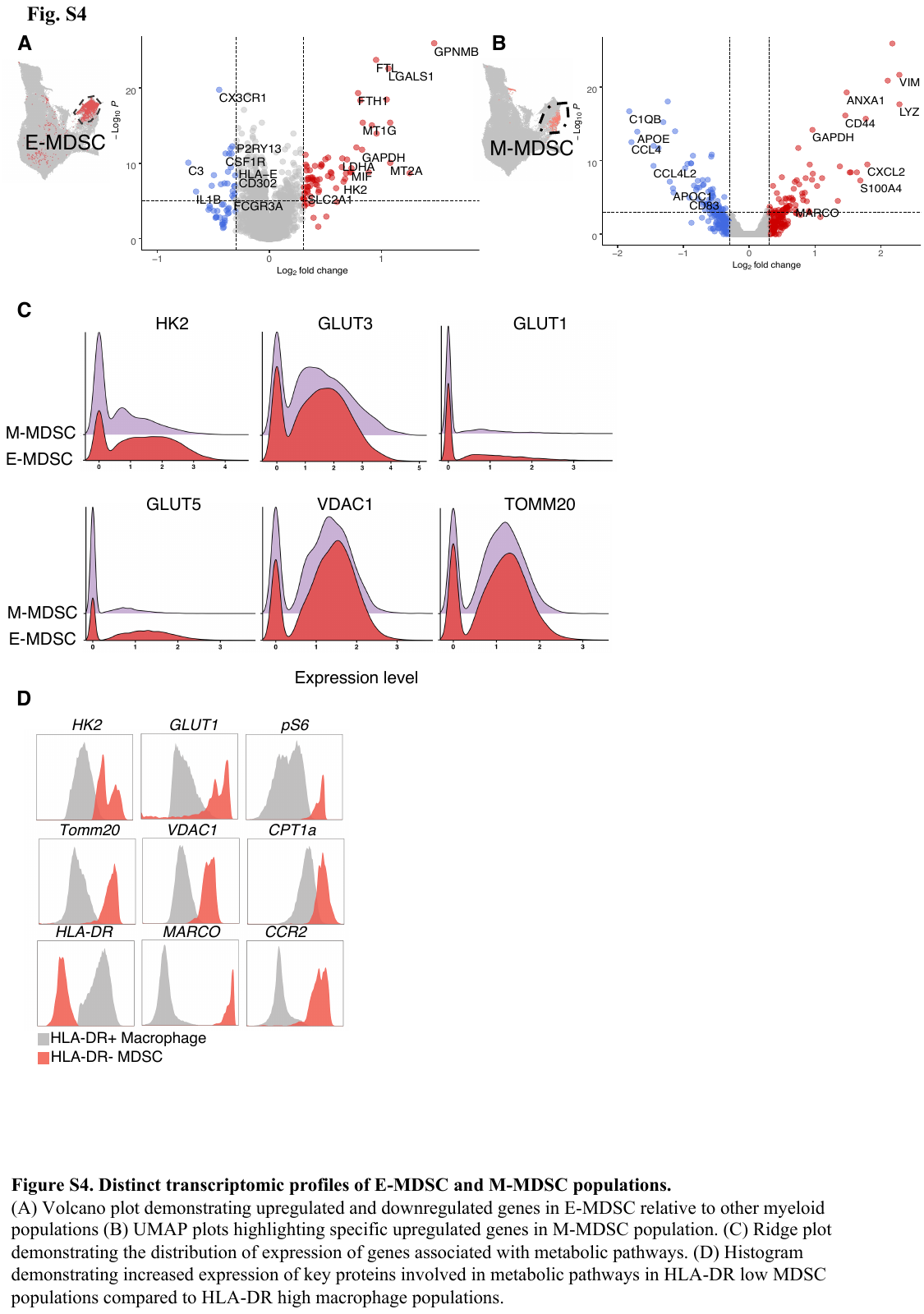

Fig. S5. Distinct transcriptomic profiles of E-MDSC and M-MDSC populations.

(A) Volcano plot demonstrating upregulated and downregulated genes in E-MDSC relative to other myeloid populations (B) Volcano plot demonstrating upregulated and downregulated genes in M-MDSC relative to other myeloid populations. (C) Ridge plot demonstrating the distribution of expression of genes associated with metabolic pathways. (D) Histogram demonstrating increased expression of key proteins involved in metabolic pathways in HLA-DR low MDSC populations compared to HLA-DR high macrophage populations.

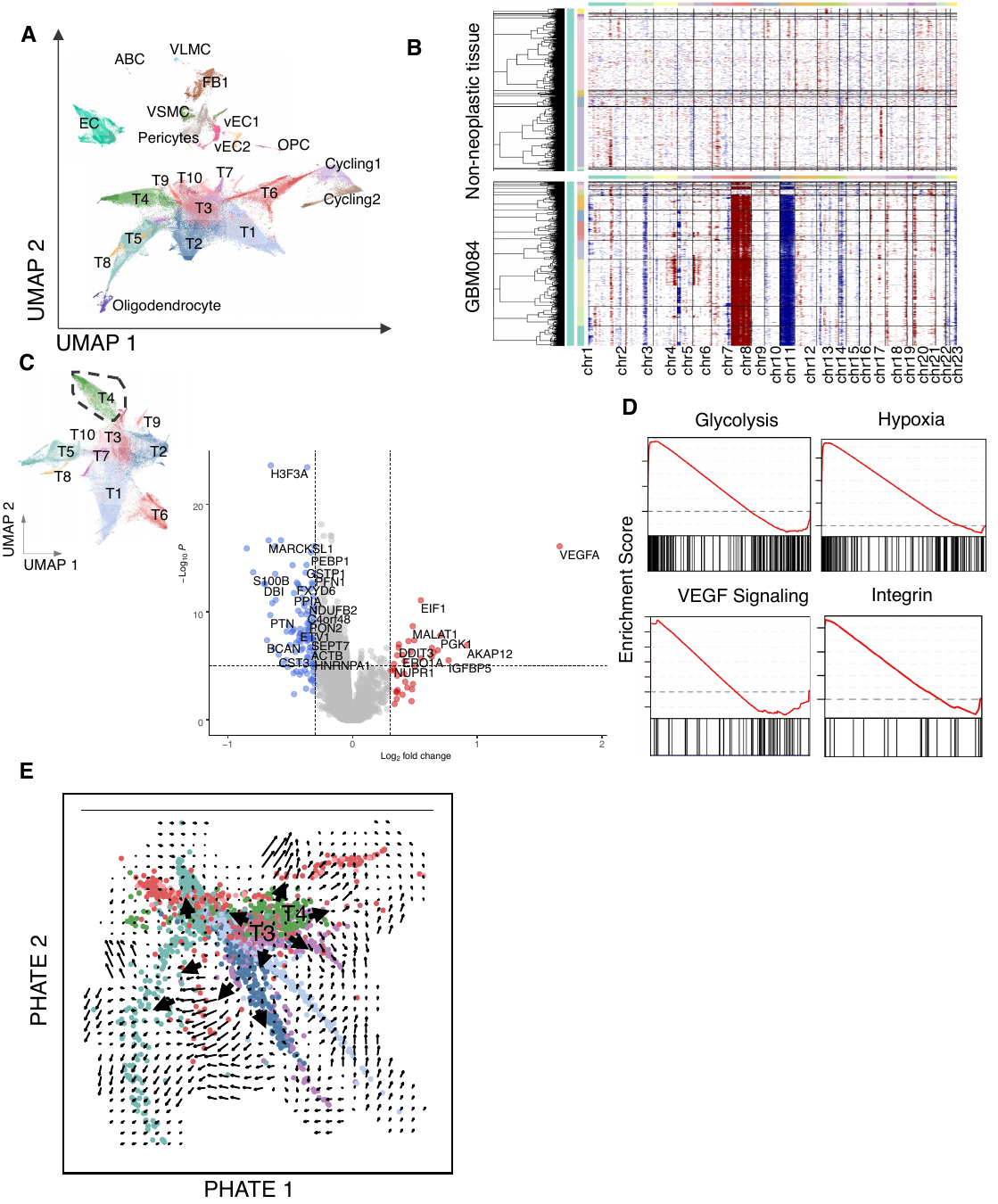

Fig. S6. Single cell atlas of CD45- cells in gliomas

(A) UMAP plot of CD45- cells in gliomas demonstrating 12 clusters of neoplastic cells and 12 clusters of stromal cells. (B) Heatmap demonstrating large-scale CNVs for individual cells (rows) across each cluster of CD45- cells from a representative tumor patient (left, GBM084), and representative non malignant tissue (right, GBM076) inferred based on the average expression of 100 genes surrounding each chromosomal position (column). Malignant cell clusters exhibit chromosome 7 gain (red) and chromosome 10 loss (blue), which are characteristic of glioblastoma. (C) Volcano plot demonstrating upregulated and downregulated genes expressed by T4 relative to other tumor populations. (D) Gene set enrichment scores of gene sets enriched in T4 demonstrating upregulation of metabolic, hypoxia, and angiogenesis pathways. (E) RNA velocities visualized on a phate map projection of tumor cells in GBM.

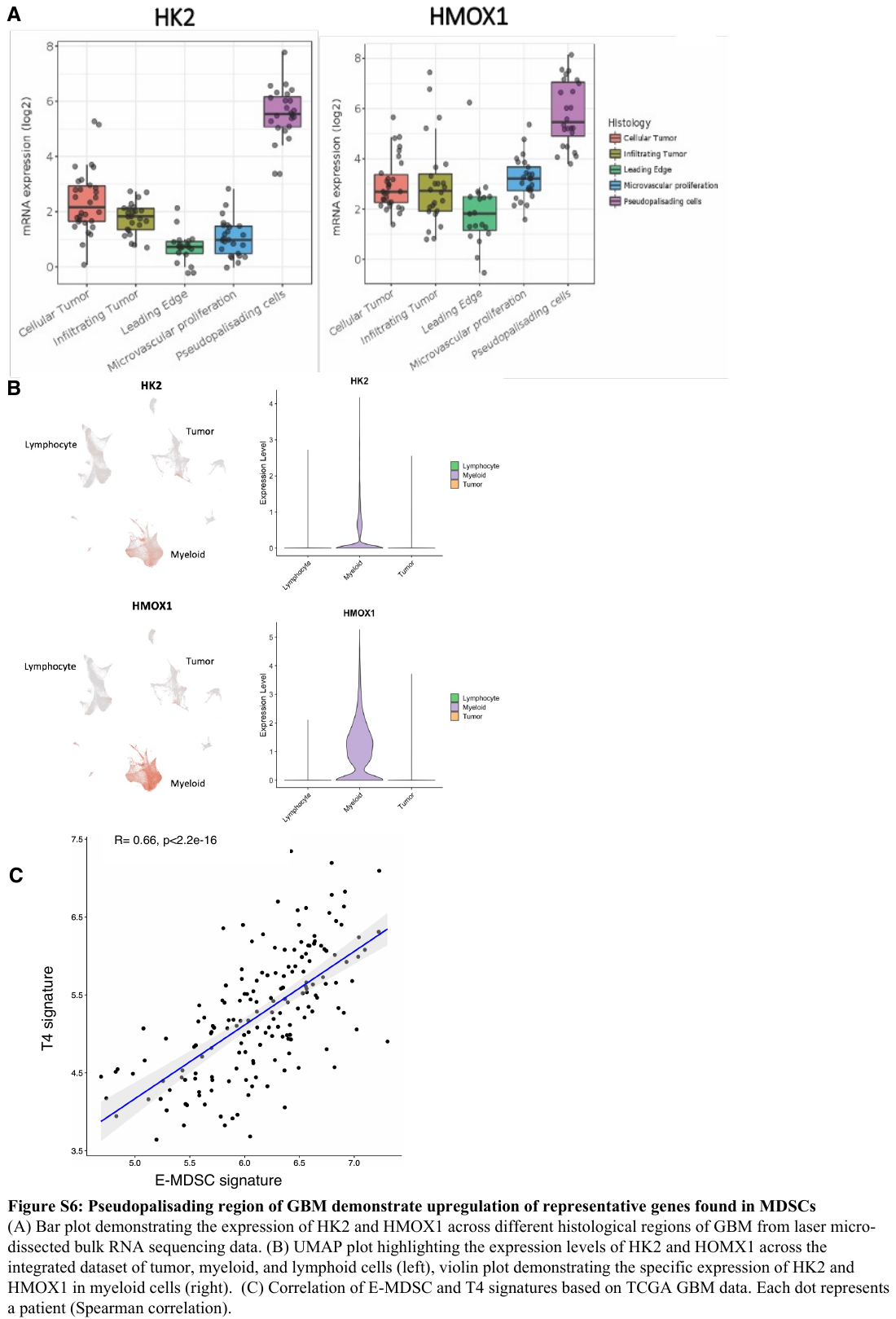

Fig. S7. Pseudopalisading region of GBM demonstrate upregulation of representative genes found in MDSCs

(A) Bar plot demonstrating the expression of HK2 and HMOX1 across different histological regions of GBM from laser micro-dissected bulk RNA sequencing data. (B) UMAP plot highlighting the expression levels of HK2 and HOMX1 across the integrated dataset of tumor, myeloid, and lymphoid cells (left), violin plot demonstrating the specific expression of HK2 and HMOX1 in myeloid cells (right). (C) Correlation of E-MDSC and T4 signatures based on TCGA GBM data. Each dot represents a patient (Spearman correlation).

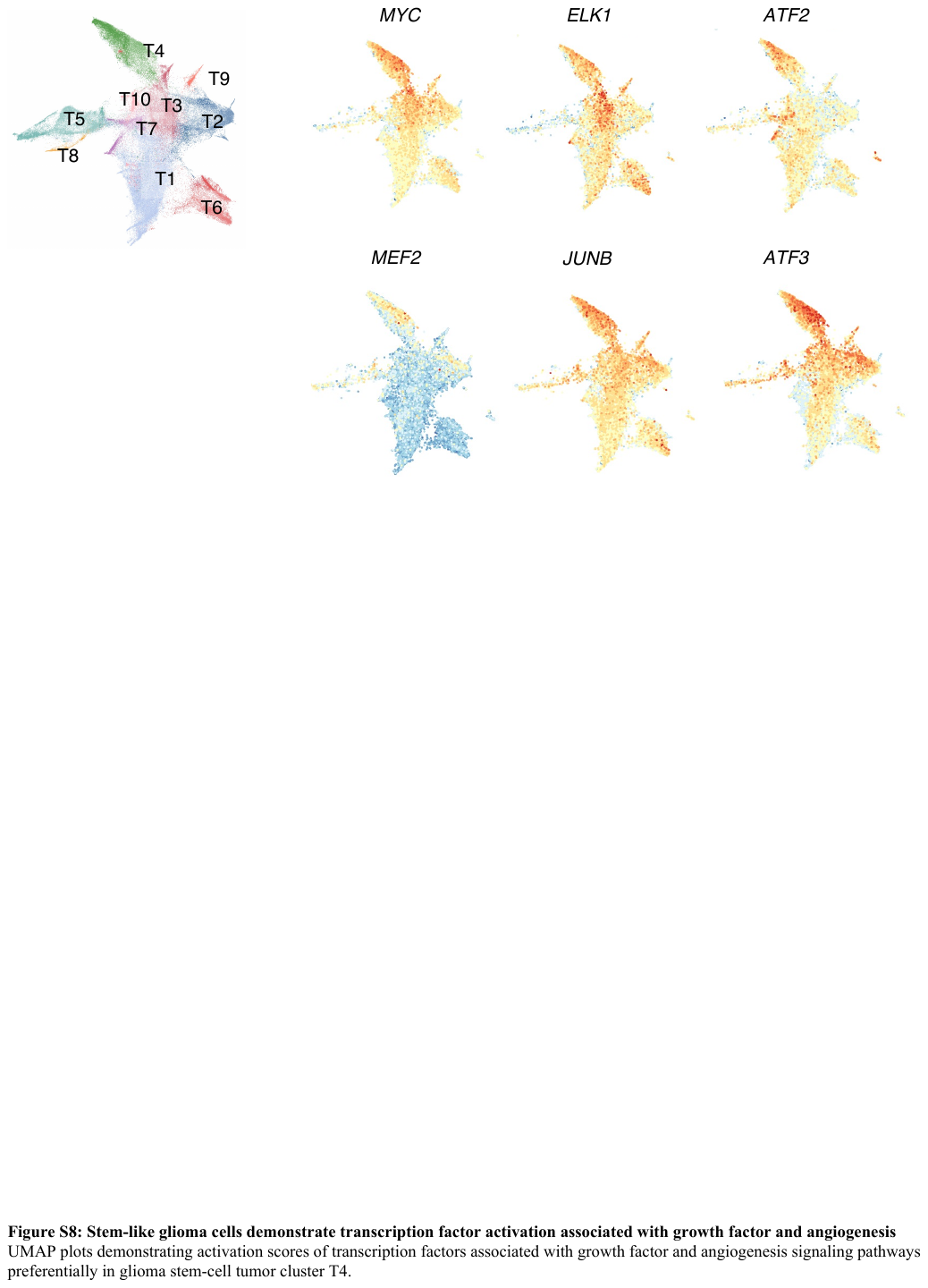

Fig. S8: Stem-like glioma cells demonstrate transcription factor activation associated with growth factor and angiogenesis

UMAP plots demonstrating activation scores of transcription factors associated with growth factor and angiogenesis signaling pathways preferentially in glioma stem-cell tumor cluster T4.

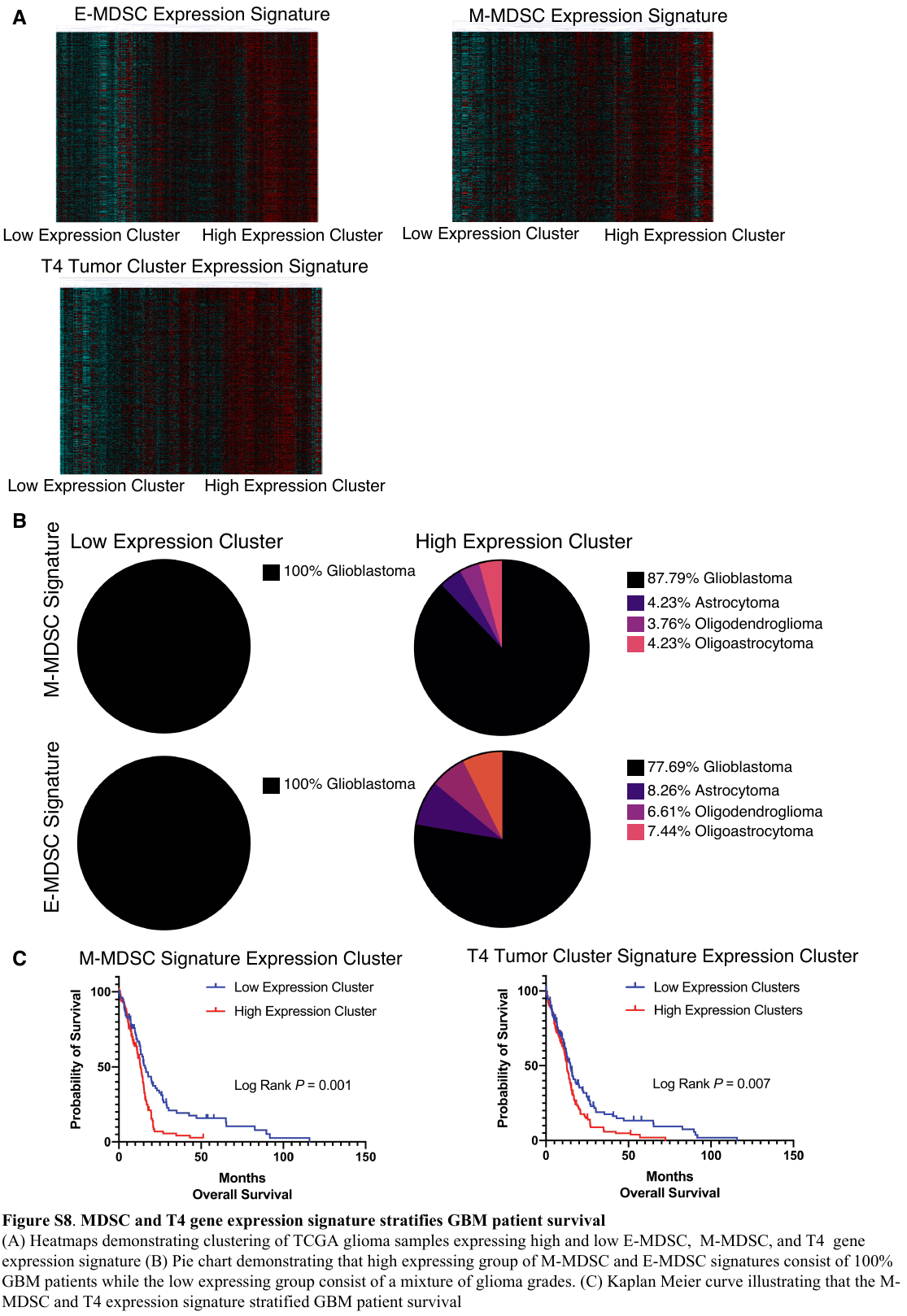

Fig. S9. MDSC and T4 gene expression signature stratifies GBM patient survival

(A) Heatmaps demonstrating clustering of TCGA glioma samples expressing high and low E-MDSC, M-MDSC, and T4 gene expression signature (B) Pie chart demonstrating that high expressing group of M-MDSC and E-MDSC signatures consist of 100% GBM patients while the low expressing group consist of a mixture of glioma grades. (C) Kaplan Meier curve illustrating that the M-MDSC and T4 expression signature stratified GBM patient survival

| **Patient ID** | **Age/Sex** | **Tumor Grade** | **Histology** | **CD45+CD3+** | **CD45+CD3-** | **CD45-** |
| --- | --- | --- | --- | --- | --- | --- |
| GBM006 | 67/F | IV | GBM | 3652 | 6841 | 3027 |
| GBM009 | 70/M | IV | GBM | 2912 | 572 | 2246 |
| GBM010 | 53/M | IV | GBM | 5829 | 1883 | 1945 |
| GBM029 | 74/F | IV | GBM | 2805 | 885 | 1332 |
| GBM030 | 66/M | IV | GBM | 5546 | 1965 | 1734 |
| GBM035 | 66/F | IV | GBM | 2415 | 1993 | 2243 |
| GBM036 | 61/F | IV | GBM | 6800 | 2087 | 8312 |
| GBM037 | 73/F | IV | GBM | 6375 | 2209 | 6310 |
| GBM043 | 63/M | IV | GBM | 7102 | 4249 | 7706 |
| GBM045 | 44/F | II | Oligo | 13018 | 3795 | 606 |
| GBM046 | 64/M | III | AA | 1735 | 3437 | 0 |
| GBM048 | 56/F | IV | GBM | 11500 | 4767 | 7220 |
| GBM049 | 71/M | IV | GBM | 11423 | 3973 | 12796 |
| GBM050 | 64/F | IV | GBM | 12818 | 7502 | 12837 |
| GBM051 | 66/M | IV | GBM | 21456 | 8879 | 7592 |
| GBM052 | 40/M | IV | GBM | 15090 | 5515 | 9924 |
| GBM054 | 64/F | IV | GBM | 9542 | 6261 | 10049 |
| GBM055 | 37/F | III | AA | 15431 | 8269 | 5925 |
| GBM056 | 78/F | IV | GBM | 12700 | 6848 | 9758 |
| GBM057 | 47/F | III | AA | 17583 | 9955 | 14802 |
| GBM059 | 38/F | II | Oligo | 7432 | 4875 | 8021 |
| GBM060 | 31/F | III | AA | 4776 | 9234 | 9922 |
| GBM064 | 28/M | IV | GBM | 12668 | 6749 | 10067 |
| GBM065 | 77/M | IV | GBM | 14104 | 5516 | 7451 |
| GBM066 | 55/M | IV | GBM | 17726 | 3563 | 8999 |
| GBM068 | 42/F | II | Oligo | 4280 | 13037 | 10448 |
| GBM069 | 66/F | II | Oligo | 11583 | 5450 | 5030 |
| GBM070 | 32/F | II | Astrocytoma | 2790 | 3993 | 10345 |
| GBM073 | 33/M | IV | GBM | 15246 | 22129 | 11674 |
| GBM074 | 64/F | II | Oligo | 17912 | 5004 | 11092 |
| GBM075 | 31/F | III | AA | 11250 | 2495 | 1880 |
| GBM076 | 22/M | Non-neoplastic | NA | 23344 | 10284 | 9529 |
| GBM077 | 33/M | Non-neoplastic | NA | 13914 | 6472 | 14279 |
| GBM081 | 24/F | II | Oligo | 6819 | 7800 | 10153 |
| GBM082 | 83/F | IV | GBM | 5596 | 11960 | 9180 |
| GBM086 | 31/F | Non-neoplastic | NA | 1761 | 3444 | 597 |
| GBM089 | 26/M | Non-neoplastic | NA | 4887 | 6787 | 5697 |
| GBM090 | 57/M | Non-neoplastic | NA | 17765 | 11471 | 19389 |
| GBM094 | 83/F | Non-neoplastic | NA | 2027 | 8045 | 5266 |

AA: Anaplastic astrocytoma, Oligo: oligodendroglioma

**Table S1.** **Patient clinical characteristic and sequencing statistics**

**Table S2. Differentially expressed genes for scRNAseq myeloid clusters.**

| **Fluorophore** | **Marker** | **Manufacturer** |
| --- | --- | --- |
| L/D | PI | Invitrogen |
| CD45 | FITC | Biolegend |
| CD3 | APC-Cy7 | BD |
| CD19 | APC-Cy7 | BD |
| CD56 | APC-Cy7 | BD |
| HLA-DR | BB700 | BD |
| CD33 | BV785 | Biolegend |
| CD14 | BV605 | Biolegend |
| CD15 | PE-Cy7 | Biolegend |
| CD16 | BV711 | Biolegend |

**Table S3. Immunofluorescence panel for sorting of MDSC subsets**

| **Fluorophore** | **Marker** | **Manufacturer** |
| --- | --- | --- |
| L/D | Zombie NIR | Invitrogen |
| CD45 | Spark NIR | Biolegend |
| CD11c | BV480 | Biolegend |
| CD3 | APC-Cy7 | Biolegend |
| CD19 | APC-Cy7 | Biolegend |
| CD56 | APC-Cy7 | Biolegend |
| HLA-DR | BV750 | Biolegend |
| CD33 | BV570 | Biolegend |
| CD14 | BV605 | Biolegend |
| CD15 | PE-Cy7 | Biolegend |
| CD16 | BV786 | Biolegend |
| CCR2 | BV510 | Biolegend |
| LOX1 | PE | Biolegend |
| CD163 | BV650 | Biolegend |
| CD206 | BV711 | Biolegend |
| MARCO | APC | eBioscience |
| HK2 | Alexa 680 | Abcam |
| VDAC1 | AF532 | Abcam |
| Tomm20 | AF405 | Abcam |
| GLUT1 | AF647 | Abcam |
| CPT1a | AF488 | Abcam |
| pS6 | AF594 | Abcam |

**Table S4. Immunofluorescence panel for the protein validation of MDSCs**

**Table S5. Ligand-receptor interaction pairs used to perform cluster interaction analysis**

**Table S6. Gene sets utilized from Molecular Signatures Database**
